# A field method to optimize dried blood spot sampling for mercury biomonitoring

**DOI:** 10.64898/2026.05.27.727713

**Authors:** Christopher J. Sayers, Liz Huamani Valdivia, Cristina P. Siguas Gonzales, Jessica N. Pisconte, Claudia M. Vega, Helen Yurek, Kevin Regan, Evan Adams, Noe R. Huaraca-Charca, Reynold Cal, Stevan Reneau, Wilber Martínez, Gilroy Welch, Kayla S. Hartwell, David C. Evers, Luis E. Fernandez, Morgan W. Tingley

## Abstract

1. Heavy metals are pervasive environmental contaminants that can impair the health of organisms globally. As the largest anthropogenic source of the potent neurotoxin mercury (Hg), gold mining has amplified these threats throughout the tropics. Consequently, there is a mounting need to monitor Hg contamination of the richest biological communities on Earth. Venous whole blood provides a reliable, nonlethal measurement of recent dietary and site-derived contamination, but collecting and cold-storing samples can be impractical in field conditions.
2. To overcome these challenges, we developed and evaluated a method to assay Hg exposure in vascular organisms by measuring the volume of dried blood spots (DBS) in the field, which can be stored at ambient temperatures until analysis. We explored the method’s precision and accuracy in estimating whole blood Hg concentrations by collecting paired whole blood and DBS aliquots from birds (*n* = 527 individuals, 140 species) along a trophic gradient (i.e., granivores to piscivores) in Belize and Peru.
3. Using a Bayesian linear mixed-effects model, we found a highly precise and unbiased relationship between DBS and whole blood total Hg concentrations that was centered at perfect unity (*R*^2^ = 0.99; *β* = 1.00 ± 0.03; 95% CrI: 0.95–1.05). Agreement between individual paired aliquots was more variable, in which approximately 12% of DBS containing at least 1 ng THg differed from whole blood by more than ±20%. However, DBS accuracy increased at higher THg concentrations, suggesting that disagreement at low concentrations is an expected consequence of higher measurement error near the analytical limit of detection of our instruments.
4. Compared to whole-blood collection and analysis workflows, DBS offer substantial logistical advantages by eliminating cold-chain dependence and reducing transport burden, laboratory handling time, and overall operational costs. Consequently, volume-measured DBS provide a practical and highly reliable alternative for monitoring Hg contamination in both humans and wildlife, particularly for ecological and population-level applications in remote and resource-limited environments.

## Introduction

Pollution of the environment by heavy metals, pesticides, microplastics, and excess nutrients presents a perennial threat to biodiversity and human health worldwide (Mueller et al., 2022). Mercury (Hg) is one example of a ubiquitous heavy metal contaminant that moves through and persists within global ecosystems for decades after emission (Gworek et al., 2020). Elemental Hg (Hg^0^) from natural and anthropogenic sources can be converted by microbes into methylmercury (MeHg), an organic form that readily biomagnifies in aquatic and terrestrial food webs and bioaccumulates to harmful concentrations in top predators (Evers et al., 2024). As a potent neurotoxin, MeHg produces a suite of deleterious effects in vertebrates, including oxidative stress, impaired endocrine, immune, and neurological function, as well as changes to behavior, reproduction, growth, and longevity (Díez, 2009; Whitney & Cristol, 2017). Artisanal and small-scale gold mining (ASGM)—the largest anthropogenic source of Hg pollution—has expanded across the global tropics in response to surging gold prices, amplifying pollution risks in the most biodiverse regions on Earth (Dethier et al., 2023; WGC, 2026). In accordance with the Minamata Convention on Mercury, a global treaty ratified by over 145 nations to protect human health and the environment from the adverse effects of Hg (UN, 2013), there is now a mounting need to monitor Hg exposure in organisms (i.e., biomonitoring) in tropical regions. However, current tools and sampling strategies may limit our ability to efficiently generate information to understand the distribution, sources, and impacts of ongoing contamination throughout the tropics.

Sampling venous whole blood from non-migratory organisms is considered the current “gold standard” for biomonitoring, as it provides a reliable, nonlethal measurement of recent dietary and site-derived Hg exposure (Evers, 2018; Evers et al., 2021). However, sampling whole blood to assess Hg contamination is both expensive and logistically challenging. Whole blood samples need to be properly sealed and stored below freezing (typically at –20°C) to minimize changes in Hg concentrations via freeze-thaw cycles, loss of water weight, external contamination, and interconversions of Hg species (Horvat & Byrne, 1992; Sommer et al., 2016; Varian-Ramos et al., 2011). Consequently, sampling whole blood in the field requires expensive, cumbersome equipment to temporarily insulate and protect blood-collection receptacles. These items can be difficult to maneuver (e.g., large coolers) and require careful handling to avoid breakage (e.g., glass capillary tubes). In remote field sites, resources needed to transport and store whole blood may preclude blood sampling as a feasible sampling methodology, and thus knowledge acquisition of biotic Hg contamination.

Dried blood spots (DBS) present a potential methodological solution for the logistical complications of collecting whole blood for Hg biomonitoring, as well as for international transport of samples. For over a century, DBS have been used as a blood storage method in human medicine (Bang, 1913; Guthrie & Susi, 1963) and pharmacology (Henderson et al., 1997), but have only recently been applied to veterinary science and toxicology (Lehner et al., 2013). To that end, biomonitoring researchers have begun employing DBS to assess Hg exposure in humans (Basu et al., 2017; Chaudhuri et al., 2009; González-Rubio et al., 2023; Nyanza et al., 2019; Santa-Rios et al., 2021), fish (Barst et al., 2020), and birds (Perkins & Basu, 2018; Shlosberg et al., 2011, 2012). DBS stored on filter paper are lightweight, portable, and circumvent whole blood transportation and storage requirements since samples can be stored at ambient temperatures for heavy metal analyses (Basu et al., 2017; Chaudhuri et al., 2009; Perkins & Basu, 2018). Concentrations of Hg in DBS are also resistant to heat sterilization, such as required upon importation by some countries to reduce disease transmission (Paul, 2005; Perkins & Basu, 2018). Depending on the supplier, filter paper cards (e.g., Whatman 903™ Proteinsaver Cards; Cytiva, USA) can also be less expensive than equivalent equipment to store whole blood, especially after eliminating the costs of refrigeration; although, some manufacturers and researchers recommend storing DBS at –20°C in a plastic bag with silica gel desiccant and humidity indicator paper to standardize blood quality and humidity (Shlosberg et al., 2011, 2012; Sjöholm et al., 2007)

Despite the growing popularity of DBS in toxicology research, there is persistent uncertainty surrounding how comparable DBS samples are to those of whole blood (Stove et al., 2012). Several studies have expressed concerns about DBS precision and accuracy related to both the internal element contamination of filter paper cards (Chaudhuri et al., 2009; Funk et al., 2013, 2015; Langer et al., 2010) and difficulties in standardizing the size of individual samples (Perkins & Basu, 2018). As a result, standardizing the volume of DBS samples (e.g., pipetting 40–50 µL onto filter paper; Barst et al., 2020; González-Rubio et al., 2023; Lehner et al., 2013; Shlosberg et al., 2012) has been recommended because using a fixed volume bypasses multiple challenges, including estimating DBS volume based on total spot area (Funk et al., 2013, 2015; Perkins & Basu, 2018), the loss of water weight when measuring sample mass (Varian-Ramos et al., 2011), and achieving the limit of quantification during laboratory analysis (Lehner et al., 2013). However, using a fixed volume as large as 50 µL not only can increase research costs—as it often requires a micropipette and disposable plastic tips (Barst et al., 2020; Lehner et al., 2013; Shlosberg et al., 2012)—but also can be impractical for field studies that sample organisms with small veins and total blood volumes (e.g., small songbirds). A key knowledge gap thus remains: can we achieve high precision and accuracy of DBS Hg concentrations through the use of small and variably sized DBS in the field? Such a method would allow the further expansion of low-cost biomonitoring opportunities throughout remote tropical locations where current need is greatest.

Given the shortcomings of whole blood sampling and current uncertainties of DBS, we sought to develop and validate a blood sampling method that is affordable, robust to environmental conditions, and suitable for small organisms. We accomplish this by measuring the volume of blood prior to DBS application in the field. Birds provide an ideal system for testing this method, as they are among the most well-studied and cost-effective taxonomic groups for monitoring environmental Hg pollution (Ackerman et al., 2016; Evers et al., 2024; Sayers et al., 2021, 2023). To evaluate our method’s replicability, we sampled birds along a trophic gradient (e.g., granivores to piscivores) in two locations (Belize and Peru) to evaluate a broad range of Hg concentrations and field conditions.

## Methods

### Study area & bird capture

We captured, nonlethally processed, and released resident and migratory bird species while performing ground-level mist-net surveys at 15 sites across central Belize and Madre de Dios, Peru between 2021–2024. To accentuate the potential scale of avian Hg bioaccumulation, we placed study sites along a gradient of anthropogenic habitat disturbance and expected Hg emission that we determined *a priori*. We surveyed artisanal gold mining operations that were active at the time of sampling or had last been active in the preceding 5 years, as well as conservation reserves without direct mining impacts. Collectively, these sites represent a variety of tropical lowland habitats (≤ 600 m in elevation; Holdridge, 1967), including upland broadleaf evergreen forest (*terra firme*), floodplain broadleaf evergreen forest (*várzea*), pine savannah, and seasonally wet grasslands (Stotz et al., 1996).

### Blood sampling

During mist-net surveys, we nonlethally collected blood samples from 527 individual birds representing 140 species following tissue collection and storage protocols provided by the Biodiversity Research Institute (Evers et al., 2021). Whenever feasible, we collected a full microhematocrit heparinized capillary tube (75 µL) of whole blood from the cutaneous ulnar vein using 27 gauge × 12.7 mm disposable hypodermic needles (BD, USA), but regularly retained smaller blood volumes when bird body mass or environmental conditions limited further collection (minimum = 5 µL). No sample exceeded the recommended limit of 1% of body mass or 10% of their total blood volume (e.g., a 7.5 g bird = 75 µL maximum blood sample permitted; Fair et al., 2010; McGuill & Rowan, 1989). We assumed that a full capillary tube represented 75 µL of blood based on the internal dimensions (75 mm × 1.15 mm) listed on the product website (Fisher Scientific, USA).

To compare Hg concentrations stored as whole blood versus DBS, we separated blood samples into two aliquots. We transferred approximately half of the sample volume onto Whatman 903™ Proteinsaver Cards (Cytiva, USA) via capillary action and stored the remaining volume as whole blood. To best estimate Hg concentrations of DBS, we quantified the volume (µL) of each spot by measuring the distance the blood occupied in the capillary tube to the nearest 0.5 mm before and after the creation of the DBS using an analog ruler or calipers, and then multiplying this value by the volume:length ratio specific to the capillary tube model (i.e., 75 µL / 75 mm = 1 µL mm^−1^; Eq. 1; Fig. 1).

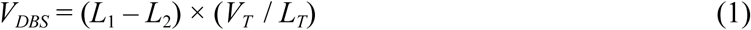

where *V_DBS_*is the volume of the DBS (µL), *L*_1_ is the length of the the whole blood sample in the the capillary tube before DBS creation (mm), *L*_2_ is the length of the whole blood sample in capillary tube after DBS creation (mm), *V_T_* is the total volume of the capillary tube (µL), and *L_T_*is the total length of the capillary tube (mm).

**Figure 1.**
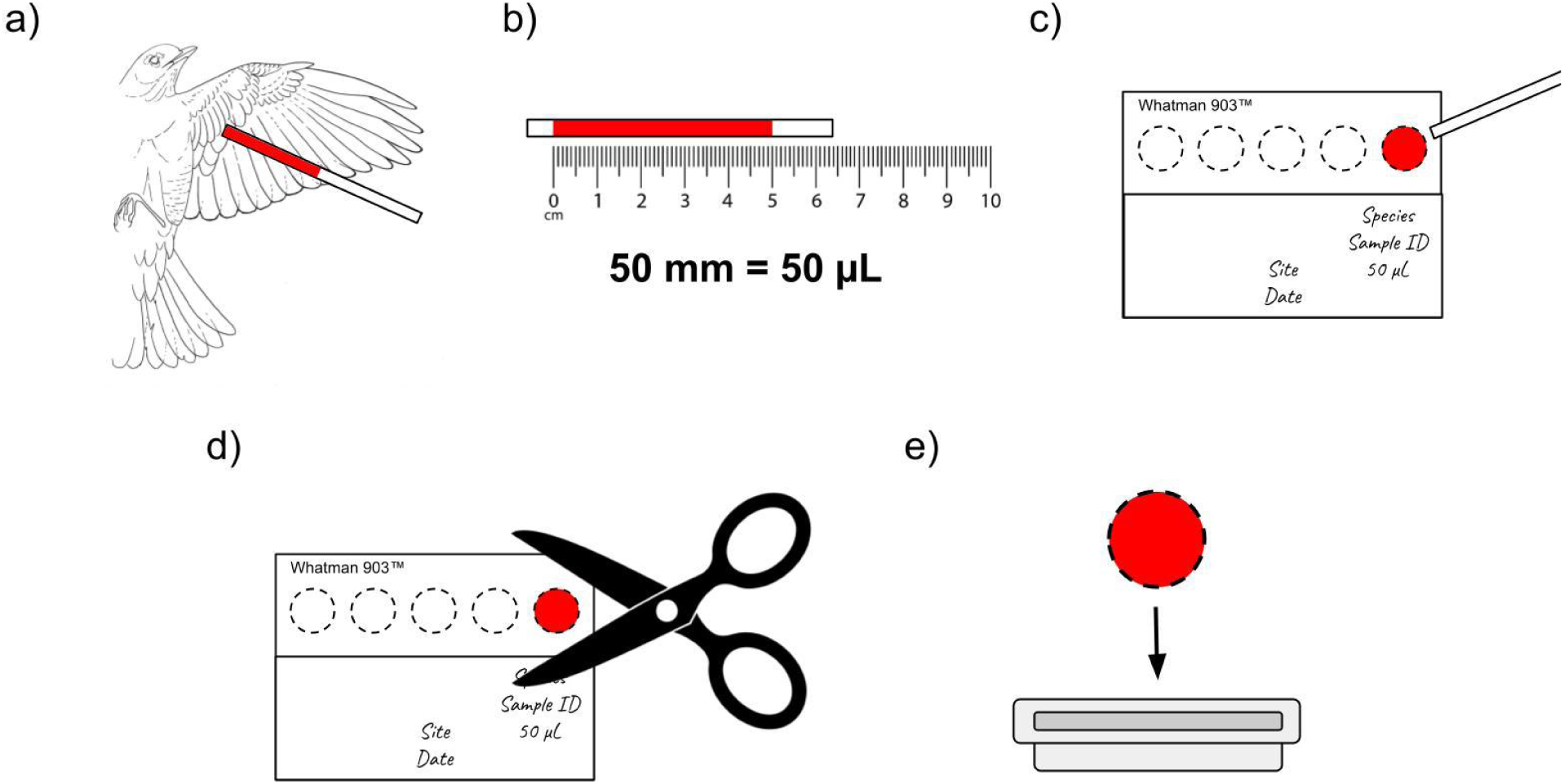
Illustration of recommended methodology: (a) extract whole blood from live bird via brachial venipuncture using a heparinized capillary tube; (b) measure length (mm) of blood sample in capillary tube using analog ruler and convert to volume (µL), which serves as the denominator in the concentration; (c) “draw” dried blood spot (DBS) on Whatman 903™ Proteinsaver Card using capillary action and label with appropriate metadata; (d) cut DBS out of card with acid-washed stainless steel scissors; and (e) place DBS into weighboat for direct mercury analysis.

We sealed capillary tubes for whole blood analysis at both ends using Hemato-Seal™ (Fisher Scientific, USA), placed tubes into a plastic 6 mL VACUETTE™ (BD, USA), and stored samples in a cooler with ice packs. We transferred whole blood samples into a freezer within 8 h of collection, where they were stored at –20°C until laboratory analysis. To freeze samples at remote field sites, we utilized an ARB 37 L portable fridge-freezer (ARB, USA) powered by a Honda EU22i portable inverter generator (Honda Motor Company, Japan). In contrast, we stored all DBS samples in plastic Ziploc™ bags at ambient temperatures without silica gel desiccant until laboratory analysis.

### Laboratory analysis

We analyzed whole blood and DBS samples for total Hg (THg) using thermal decomposition and atomic absorption spectrophotometry with either a Milestone DMA-80 direct Hg analyzer at the Laboratorio de Mercurio y Química Ambiental (Puerto Maldonado, Madre de Dios, Peru; 62% of samples) or a Nippon MA-3000 at the Biodiversity Research Institute Toxicology Lab (Portland, Maine, USA; 38% of samples). We followed United States Environmental Protection Agency Method 7473 (SW-846; USEPA, 1998) during both procedures and assumed nearly all THg (> 95%) detected in blood samples was MeHg (Edmonds et al., 2010; Rimmer et al., 2005).

To load whole blood samples into the analyzer, we removed the wax sealant from both ends of the capillary tube using a hypodermic needle, ejected the blood from the tube into a pre-tared nickel or ceramic weighing boat using a silicone bulb (Globe Scientific, USA), and measured the mass of each sample to the nearest 0.001 mg on a OHAUS Explorer™ Semi-Micro EX225D analytical balance (OHAUS, USA). For DBS, we cut out blood samples from filter paper cards by hand with stainless steel scissors washed with 2% hydrochloric acid, cutting along the dotted line to ensure the total dried blood volume was present in each cut-out. We folded paper circles in half to better secure them in the weighing boat before placing them in the analyzer. Measuring the mass of DBS samples in the laboratory was unnecessary because we already measured the sample volume in the field. We express whole blood THg concentrations as ng Hg/mg blood wet weight and DBS THg concentrations as ng Hg/µL blood, where both ratios equate to parts per million (ppm).

To ensure analytical precision and accuracy, we used quality control methods and certified reference materials, including DOLT-5, IAEA-436A, IAEA-476, ERM-CE464, and CRM-13, which all had recovery errors within ±10%. We then analyzed blank circles from cards exposed to either laboratory or field conditions to quantify potential pre-existing internal Hg contamination of filter paper cards and external contamination of DBS samples in the field (Funk et al., 2013, 2015). Using the mass of blank samples to determine their concentration in ng THg/mg, we analyzed one blank circle from five separate cards across five different lot numbers that did not leave the laboratory (*n* = 25) and one blank circle from five separate cards exposed to field conditions at eight sampling sites (*n* = 40). We excluded any blood or blank sample that registered below the lower detection limits of our instruments from the final database (Milestone DMA-80: < 0.01 ng THg, Nippon MA-3000: < 0.001 ng THg; *n* = 10 blood samples, 21 blanks), since correcting these detections could bias a linear regression (Shoari & Dubé, 2018).

### Statistical analysis

To assess the overall precision and accuracy of DBS in estimating whole blood THg concentrations, we quantified the relationship between paired blood samples using a linear mixed-effects model in a Bayesian hierarchical framework that simultaneously accounts for measurement error of both DBS and whole blood contamination across instruments (R package “R2jags”; Su & Yajima, 2024). This model is an error-in-variables regression, in which we treated the true response as a latent variable and estimated separate variance parameters for each Hg analyzer. The result is a model analogous to orthogonal regression in a frequentist approach (Fang et al., 2017; Reilly & Patino-Lea, 2012), but with the added benefit of accounting for pseudoreplication via random effects. We arbitrarily assigned DBS THg concentrations (ng/µL) as the response variable and whole blood THg concentrations (ng/mg wet weight) as the only fixed effect, and included sampling site (*n* = 15) and species (*n* = 140) as random slopes of whole blood THg. We fixed the global intercept at zero to reflect the expectation that zero Hg in DBS corresponds to zero Hg in whole blood, and estimated all other parameters using vague priors (Eq. 2). The model included three chains that ran for 55,000 iterations, including a burn-in of 5,000 and 50-fold thinning. We present all posterior estimates as model-derived predictions with accompanying 95% credible intervals. The formal model description is as follows:

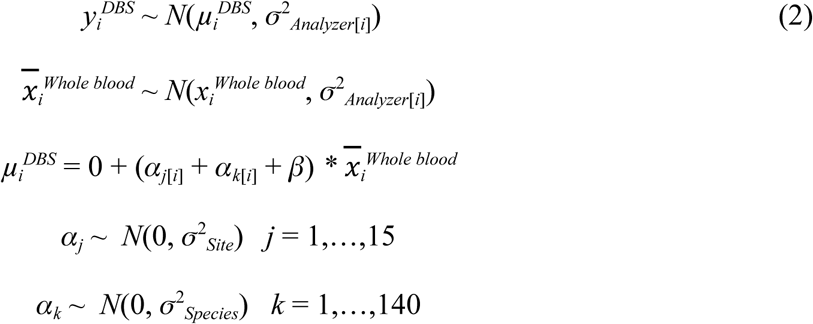

where 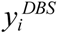 and 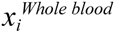 are the observed THg concentrations for individual *i* using DBS or whole blood, respectively; 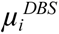 and 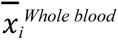 are the latent true THg concentrations for individual *i*; *β* is the overallslope relating whole blood THg to DBS; *⍺_j_*_[*i*]_ is the additive random effect on the slopefor site *j*; *⍺_k_*_[*i*]_ is the additive random effect on the slope for species *k*; and *σ*^2^ are variance parameters estimated for each Hg analyzer, as well as hierarchically for site and species.

Finally, to quantify the accuracy of individual DBS samples, we calculated the ratio of DBS to whole blood THg concentrations as a percent recovery for each paired sample (Perkins & Basu, 2018; USEPA, 1998). As measurement error is expected to increase towards the analytical limit of detection during direct Hg analysis (i.e., at very low values), we restricted our inference to paired samples above 1 ng THg (*n* = 300).

## Results

The relationship between THg concentrations of paired DBS and whole blood aliquots was both highly precise and unbiased, with a slope centered at perfect unity (*R*^2^ = 0.99; *β* = 1.00 ± 0.03 [mean ± standard deviation]; 95% CrI: 0.95–1.05; Table S1; Fig. 2). In contrast to past studies, contamination of the filter paper cards had no apparent influence on variation between paired DBS and whole blood samples, as THg concentrations were negligible across all blanks (maximum = 0.003 µg/g fw; Table S2). Furthermore, unequal measurement error between our direct Hg analyzers—due to inherent differences between their precision and detection limits—had minor contributions to the width of our credible interval (σ_Milestone DMA-80_ = 0.04 ± 0.00, σ_Nippon MA-3000_ = 0.02 ± 0.00; Table S1). Despite strong agreement across paired samples, variation between individual samples was more pronounced, with 12% of paired aliquots containing more than 1 ng THg falling outside the acceptable ±20% range of sample deviation defined by United States Environmental Protection Agency Method 7473 (USEPA, 1998; Fig. S4). However, DBS accuracy increased with more ng THg detected (Fig. S5), suggesting that disagreement at low concentrations is an expected consequence of higher measurement variation and proportional error towards the analytical limit of detection of our instruments. Given current pricing (2026), we calculated that our methodology is 15% ($0.19) less expensive than traditional whole blood sampling at $1.10/sample (Table 1), excluding additional costs to transport and freeze whole blood samples prior to Hg analysis.

**Figure 2.**
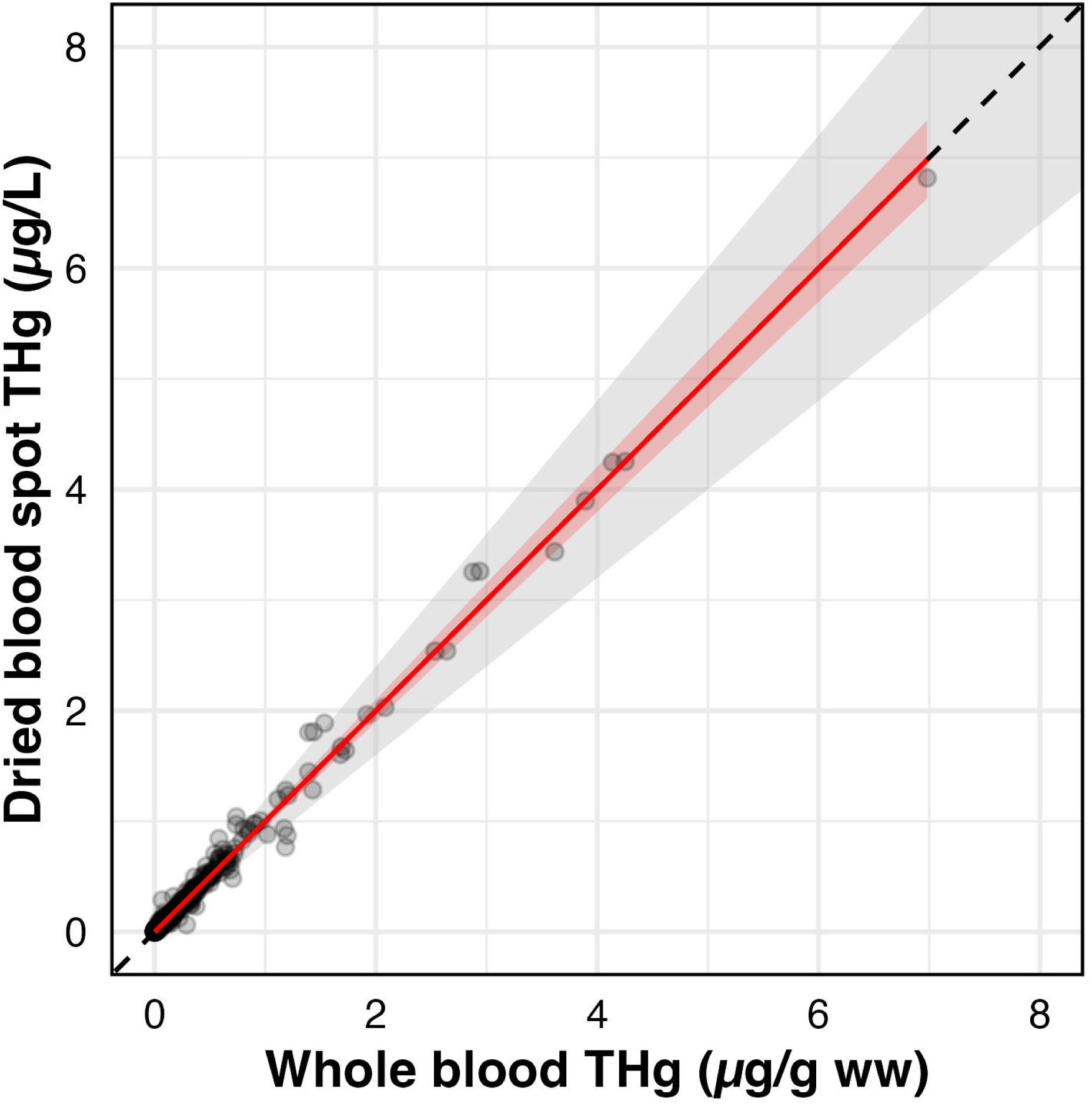
Bayesian error-in-variables linear regression of total mercury (THg) concentrations from paired dried blood spot (DBS) and whole blood samples (*n* = 527; *R*^2^ = 0.99), represented by a solid red mean line and a shaded red 95% credible envelope. The black dashed line represents a perfect 1:1 relationship between DBS and whole blood THg concentrations, while the gray shaded region represents the industry-standard acceptable range of sample deviation based on United States Environmental Protection Agency Method 7473 (±20%; USEPA, 1998).

**Table 1.**
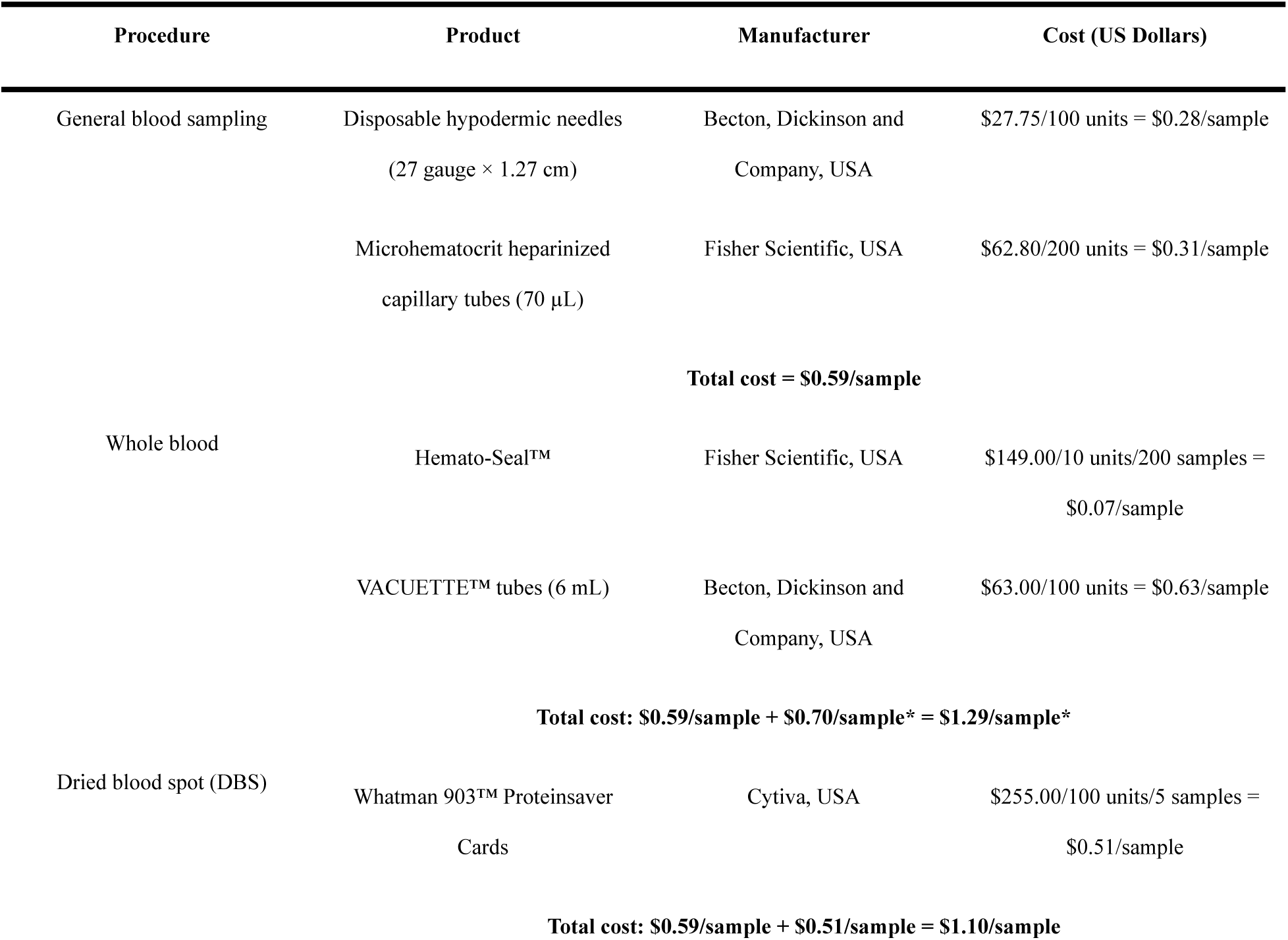
Equipment costs (USD) for collecting whole blood and dried blood spots (DBS) for Hg analysis. Other necessary consumable items (e.g., plastic bags and permanent markers) and fixed costs (e.g., handheld cooler and ice packs) were excluded for brevity. An asterisks (*) indicates that the sum does not include the cost of refrigeration, which varies based on machine energy efficiency and geography. Prices verified on Fisher Scientific (https://www.fishersci.com/) and Sigma-Aldrich and (https://www.sigmaaldrich.com/) on 28 April 2026, but subject to change.

## Discussion

In this study, we developed a simple, inexpensive blood sampling method to monitor Hg contamination in vascular organisms and evaluated its performance by sampling birds along trophic and Hg-emission gradients in central Belize and Madre de Dios, Peru. Measuring the volume of DBS in the field facilitated highly precise and unbiased estimates of whole blood THg concentrations, indicating that DBS can reliably quantify variation in Hg exposure across direct Hg analyzers, individuals, species, and sites under tropical field conditions. Although agreement between individual paired samples was more variable at low THg concentrations, recovery became more consistent as concentrations increased, suggesting that the observed disagreement was driven by higher proportional error near the analytical limit of detection rather than systematic bias in either sampling method.

### Bias & contamination of DBS

Several previous studies have reported systematic positive bias in DBS heavy metal concentrations relative to paired whole blood aliquots when measuring the mass of DBS samples, attributing the discrepancy to background contamination from filter paper cards (Chaudhuri et al., 2009; Funk et al., 2013, 2015; Langer et al., 2010). In contrast, we detected no average bias in DBS THg concentrations after directly measuring the volume of DBS in the field. Furthermore, we found no evidence of systematic internal or external contamination of filter paper cards. Blank samples exposed to both lab and field conditions had similar means and standard deviations, and no blank exceeded 0.003 µg/g fw THg (Table S1). Consequently, any potential contamination that we observed is unlikely to be biologically meaningful when describing Hg risk, since many sublethal effect thresholds for avian Hg exposure begin at concentrations several orders of magnitude higher (e.g., 0.7 ppm in the songbird, *Thryothorus ludovicianus*; Jackson et al., 2011). Together, these findings suggest that previous reports of DBS bias may partly reflect methodological limitations associated with quantifying DBS concentrations using sample mass due to the unequal distribution of Hg across individual DBS (Perkins & Basu, 2018). Our approach of directly quantifying DBS volume reduces this source of variation and provides renewed confidence for DBS as a reliable substitute for whole blood in mercury biomonitoring applications.

### Mercury recovery of DBS

To ensure consistent analytical precision during Hg analysis, United States Environmental Protection Agency Method 7473 defines an acceptable range of variation between individual Hg samples, often referred to as Hg recovery (Perkins & Basu, 2018), as ±20% deviance (USEPA, 1998). Here, we apply this threshold as a practical analytical benchmark for comparison rather than as a formal equivalence standard between DBS and whole blood sampling workflows. Although 12% of paired samples containing more than 1 ng THg exceeded this recovery range (Fig. S4), DBS accuracy improved with more ng THg detected in a sample. As expected, variation in Hg recovery was not uniform across the observed range of THg, with recovery being most variable at low THg concentrations and becoming more constrained as THg increased (Fig. S5). This pattern arises because small absolute differences between DBS and whole blood at low THg concentrations translate to large proportional error, which is then amplified by decreasing analytical precision towards the limit of detection of our instruments—increasing the likelihood that low THg samples exceed a fixed threshold. Additional sources of variation, such as measurement error when estimating DBS volume or heterogeneity in blood composition (e.g., quantity or distribution of MeHg bound to red blood cells; Berlin, 1963; Kershaw et al., 1980; Norseth & Clarkson, 1970), may further contribute to reduced DBS accuracy, especially for samples with low concentrations. Taken together, these results suggest that the lack of agreement between DBS and whole blood at low concentrations reflects expected proportional error and the analytical limitations of our instruments, rather than the poor performance of DBS themselves. Accordingly, metrics based on fixed proportional thresholds, such as Hg recovery, are inherently sensitive to non-uniform error structures (i.e., heteroscedasticity), and calibration-based approaches may provide a more reasonable assessment of method performance.

### Benefits of this methodology

Volume-estimated DBS offer several logistical, economic, and scientific advantages over both whole blood and past DBS methodologies. By eliminating cold-chain requirements, DBS simplify nearly every stage of sample handling—from field collection to laboratory analysis—while reducing overall operational costs. Compared to whole blood, DBS are less expensive to collect on a per-sample basis (Table 1) and eliminate the need for infrastructure to keep samples frozen in the field, including a portable freezer, generator, and petroleum fuel. DBS also reduce handling time and contamination risk in the laboratory, as technicians do not need to thaw frozen samples, remove clay sealant from capillary tubes, nor measure sample mass prior to analysis.

The primary innovation of our method lies in manually measuring DBS blood volume in the field, rather than obtaining a fixed volume using additional, more expensive equipment (e.g., a micropipette and many disposable plastic tips). Collecting variably sized DBS is particularly advantageous when working with small organisms, such as songbirds, since this approach can accommodate natural stochasticity in sample volume during venipuncture in the field—an option that could also safeguard animal welfare by allowing smaller blood draws proportional to a bird’s total blood volume (Fair et al., 2010; McGuill & Rowan, 1989). Beyond Hg quantification, paired DBS samples can also support a range of genetic and genomic analyses—including demographic estimation, whole genome sequencing, and blood parasite screening—due to the high DNA yield associated with nucleated avian red blood cells (Samsonova et al., 2022).

Taken together, volume-estimated DBS stand to eliminate many logistical and financial barriers associated with Hg biomonitoring, especially in remote regions with limited infrastructure. Indeed, this is only the second study to our knowledge that uses field-prepared DBS to assess Hg exposure in wild organisms (Barst et al., 2020), and a first for birds. With additional validation, the potential exists to extend this methodology to other heavy metals or blood contaminants that remain stable in DBS.

### Recommendations and future directions

Based on our findings, we suggest the following refinements to improve efficiency and reduce potential sources of error:

1. Practitioners of this method must take extreme care to avoid introducing air bubbles into capillary tubes during venipuncture, as any trapped air will bias DBS volume estimates by artificially increasing the measured length of blood within the tube.
2. Although a standard filter paper card can physically accommodate up to five 100 µL DBS samples, we caution against storing more than 2–3 samples of approximately 75 µL per card to prevent oversaturation and cross-contamination via lateral bleeding between samples (see Chaudhuri et al., 2009). Blood volumes between 10–75 µL are usually sufficient for THg analysis, however, we recommend 30–60 µL as an ideal range to exceed the lower analytical limit of quantification, minimize animal stress during venipuncture, and avoid oversaturation. As a best practice, unused blank DBS circles should be analyzed systematically per card to estimate potential external Hg contamination from the field.
3. To standardize sample volume and minimize equipment costs for animals of sufficient mass, researchers should consider using smaller capillary tubes (e.g., 50 µL models) that are easier to fill completely and thus do not need to be manually remeasured to determine blood volume (see González-Rubio et al., 2023).
4. Some prior research has recommended storing DBS at –20°C in a plastic bag with silica gel desiccant and humidity indicator paper to preserve blood quality and standardize sample moisture prior to analysis (Shlosberg et al., 2011, 2012; Sjöholm et al., 2007). Given the precision and accuracy of our results despite storing samples at ambient tropical temperatures without desiccant, and that our method does not rely on sample mass to quantify Hg concentration, we presently view these steps as superfluous and costly. Perkins & Basu (2018) also report that these storage recommendations do not have an apparent effect on DBS accuracy.
5. To standardize the amount of excess filter paper—and accompanying potential contamination—surrounding DBS samples, laboratory technicians should use a 15 mm hole punch to extract DBS instead of cutting them out with scissors.
6. Having demonstrated strong agreement between volume-estimated DBS and whole blood THg concentrations in Neotropical birds, we encourage other field researchers to evaluate our approach for other contaminants, organisms, and ecosystems.

## Conclusions

Collecting DBS by measuring the sample volume in the field offers a practical and highly reliable alternative to sampling whole blood in ecotoxicology research. DBS are more affordable, portable, and robust to field conditions than whole blood, especially when considering the financial costs of refrigeration and the logistical burden associated with keeping samples frozen in the field. Analyzing DBS also requires less labor in the laboratory, since they eliminate the need to thaw frozen samples, extract clay sealant from capillary tubes, and measure the mass of samples. Their high precision and agreement with whole blood make DBS well-suited for detecting relative differences in Hg contamination across populations and sites. However, caution may be warranted for applications that require strict agreement with individual whole blood concentrations, especially when background contamination is low. By reducing economic and logistical barriers, this method has strong potential to expand research on Hg contamination of birds and other vascular animals in remote and resource-limited environments facing growing pollution concerns, particularly the global tropics.

## Data availability

Supplementary methods and results are available in Appendix S1. All data and software curated for this research are available from the Dryad Digital Repository (future link here).

## Acknowledgements

We thank Julissa Barrios Ccoyori, Arturo Mamani Ramos, Erick Huaman Vargas, Raúl Mandujano Collantes, Teresa Ávalo Vilchez, Jorge Novoa Cova, Adamelita Quispe, Daniel Carpio, Darwin Mamani Mamani, Ruth Caviedes Ccoyuri, Jhoanna Cortegano Tapullima, Victor Ramos Ascue, Lucero Horna Ordinola, Jelicsa Peña Girón, Finola Fogarty, Julio Salvador, Larry Huacarpuma-Aguilar, and Abidas Ash for assistance in the field; Jhon Farfán, France Cabanillas, Mathieu Charette, José Purisaca Puicón, and Mariana Valqui for logistical assistance; and Inkaterra Asociación, Lago Soledad Tented Camp, Los Amigos Biological Station, Foundation for Wildlife Conservation, Tropical Education Center, Toucan Ridge Ecology & Education Society, and Monkey Bay Wildlife Sanctuary for granting site access.

## Ethics

Sampling was conducted with authorization from the Belize Forest Department (FD/WL/1/21 [19]), Servicio Nacional Forestal y de Fauna Silvestre (SERFOR; N° 17-MAD-TAM/AUT-IFS-IFL-2023-001) and Servicio Nacional de Áreas Naturales Protegidas por el Estado (SERNANP; 17-2023-SERNANP-JEF) in Peru, as well as Animal Care and Use Committees (IACUC) of University of California, Los Angeles (ARC-2017-073).

## Author contributions - CRediT author statement

Conceptualization: CJS, KR, DCE; Methodology: CJS; Validation: CJS, LHV, HY; Formal analysis: CJS, MWT; Investigation: CJS, LHV, CPSG, JNP, NRHC, RC, SR, WM, GW; Resources: LHV, CMV, HY, KR; Data Curation: CJS, LHV, HY, KR; Writing - Original Draft: CJS; Writing - Review & Editing: MWT, LHV, JNP, CMV, KR, EA, DCE, LEF; Visualization: CJS; Supervision: MWT, CMV, LEF, DCE; Project administration: CJS, JNP, LHV, CMV, KSH; Funding acquisition: DCE, LEF, MWT, CJS.

## Statement on inclusion

Our study brings together authors from several countries, including scientists that live and work in the countries where we conducted the study. All authors engaged early in the study design to ensure a diverse set of perspectives were considered. Whenever relevant, we cited literature published by scientists from the study regions, and considered work published in local languages.

## Supplementary Materials

### I. Bayesian linear regression

**Table S1.**
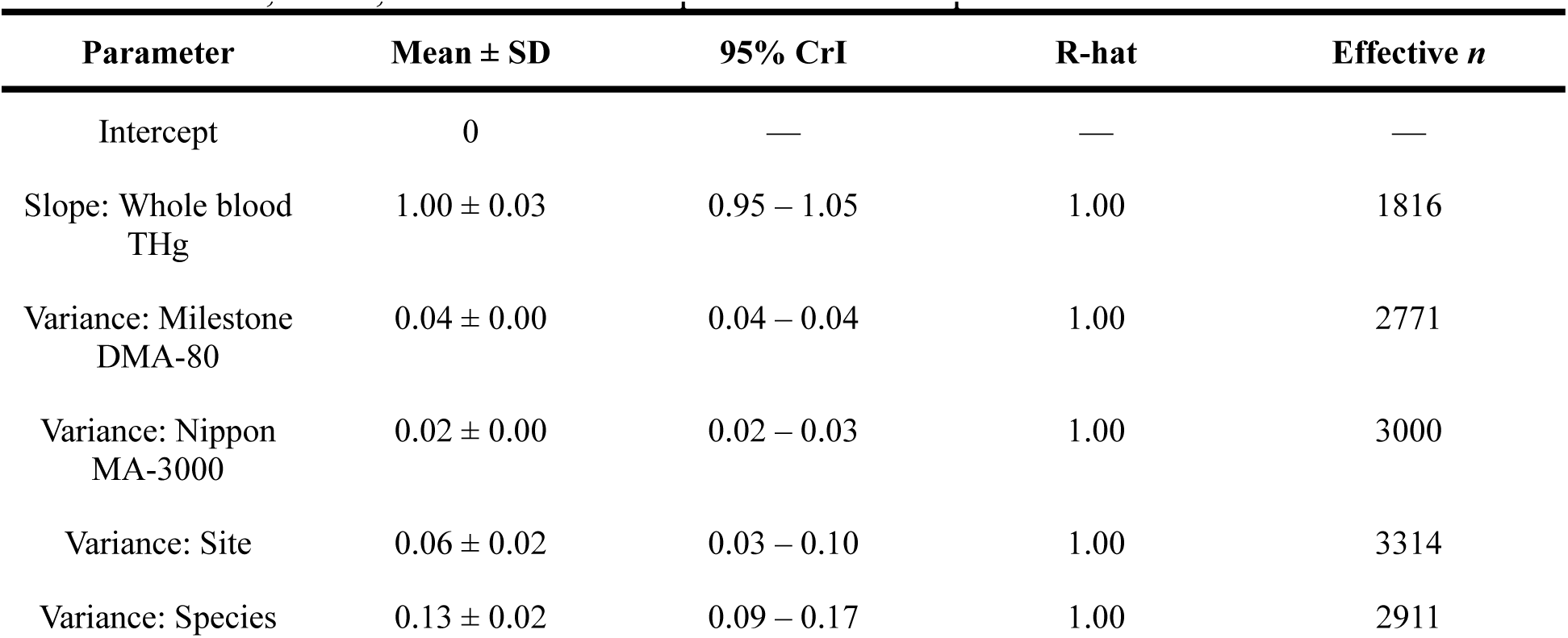
Summary of Bayesian linear model with the posterior mean ± standard deviation, 95% credible interval, R-hat, and effective sample size for each parameter.

**Figure S1.**
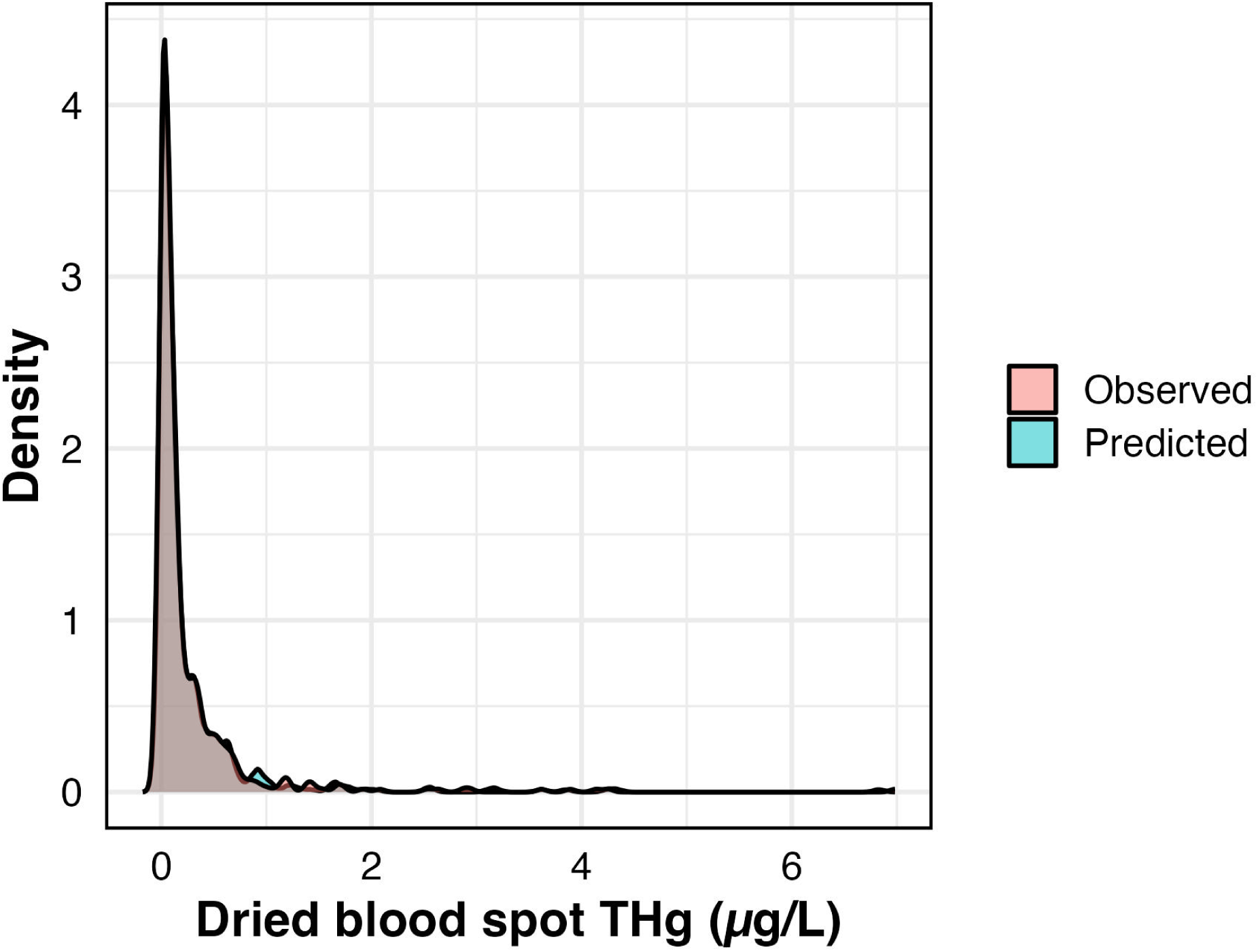
Posterior predictive check of model-derived predictions fit to the observed data.

**Figure S2.**
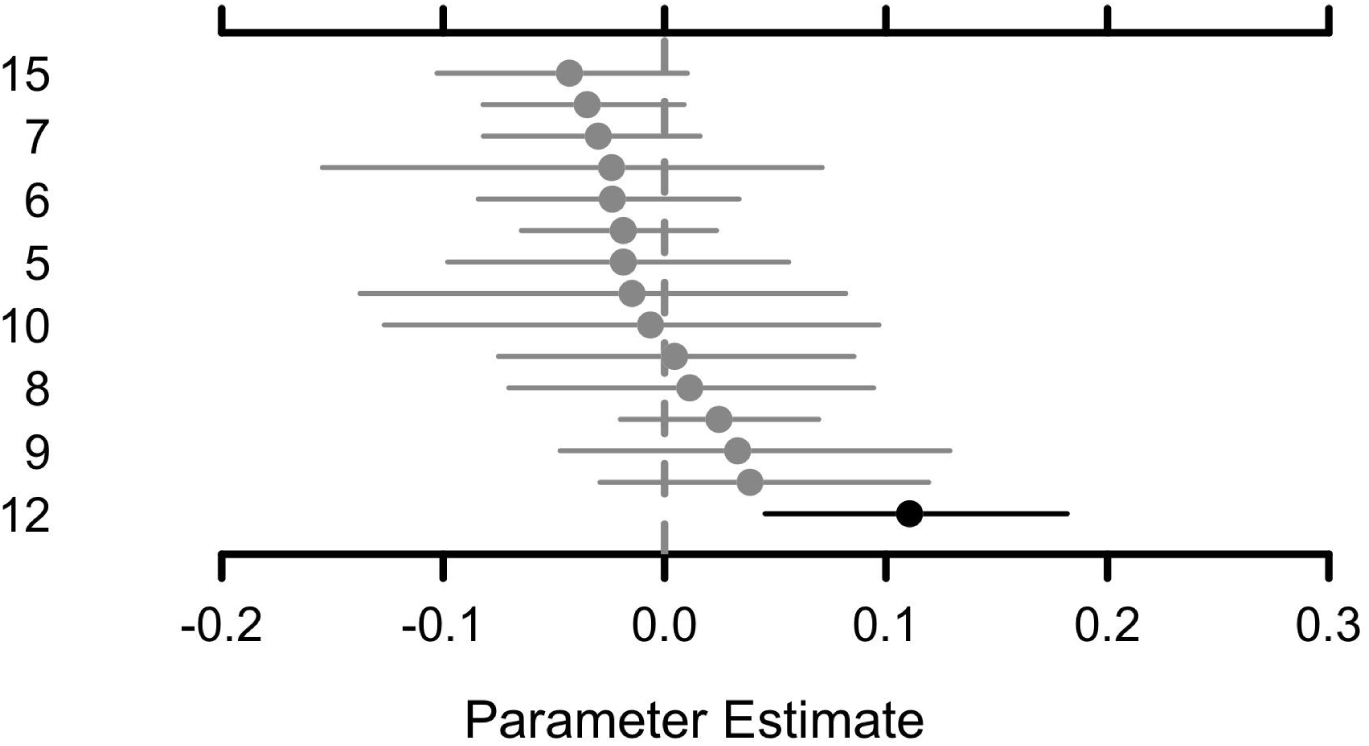
Effect plot of random site slopes for whole blood THg. Point ranges in black indicate parameters with credible intervals that do not overlap zero and are statistically significant.

**Figure S3.**
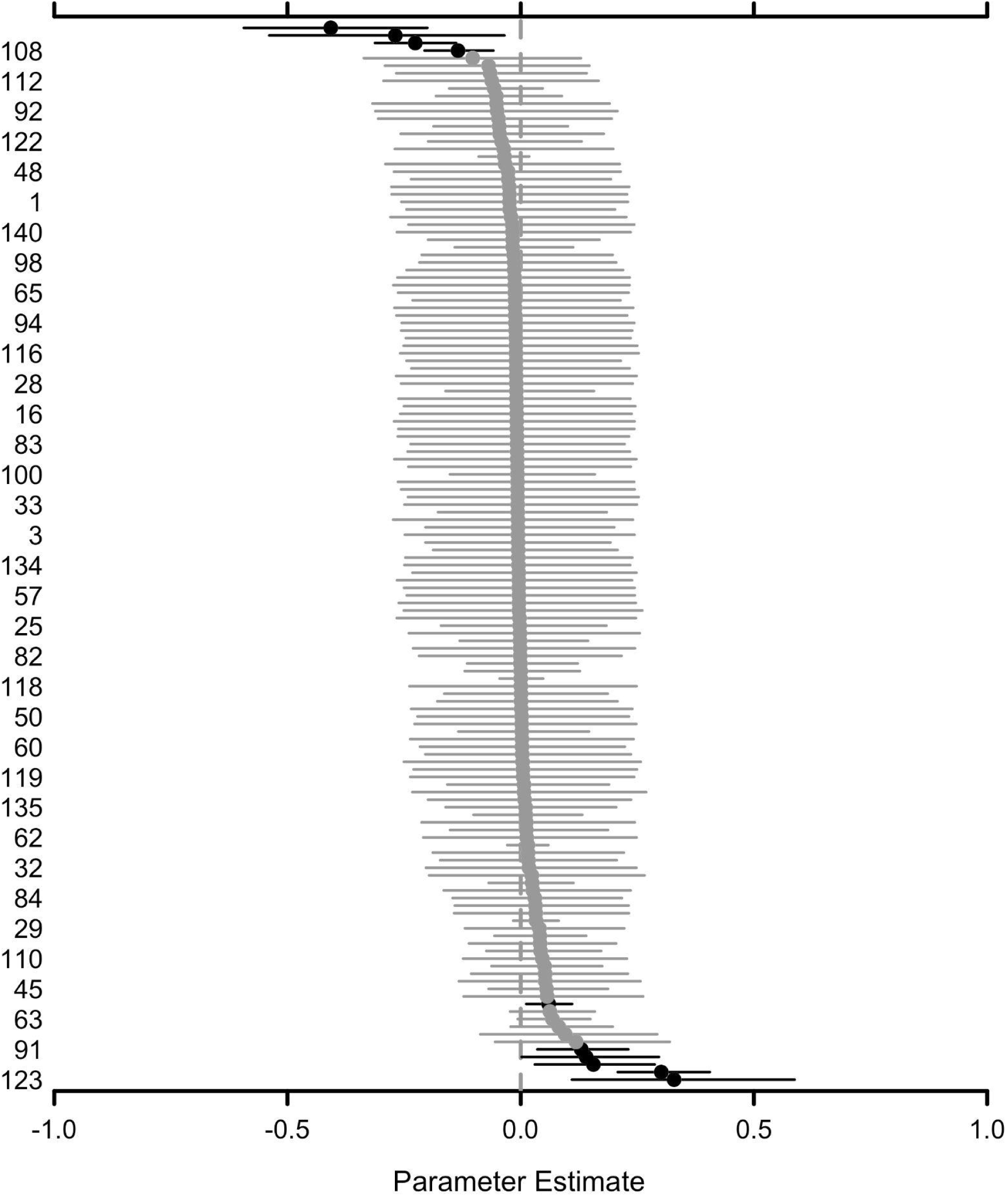
Effect plot of random species slopes for whole blood THg, in which names on the vertical axis are thinned for legibility. Point ranges in black indicate parameters with credible intervals that do not overlap zero and are statistically significant.

### II. Blanks

**Table S2.** Total mercury (THg) concentrations (µg/g fresh weight) of blank filter paper samples exposed to laboratory and field conditions. All samples that registered below the lower limit of detection (*n* < LoD) were excluded from calculations.

| Exposure regime | *N* | *n* < LoD | *n* | Mean ± SD | Range |
| --- | --- | --- | --- | --- | --- |
| Laboratory | 25 | 5 | 20 | $0.0007 \pm 0.0005$ | 0.0001 – 0.0017 |
| Field | 40 | 16 | 24 | $0.0009 \pm 0.0006$ | 0.0004 – 0.0030 |

### III. Mercury recovery

**Figure S5.**
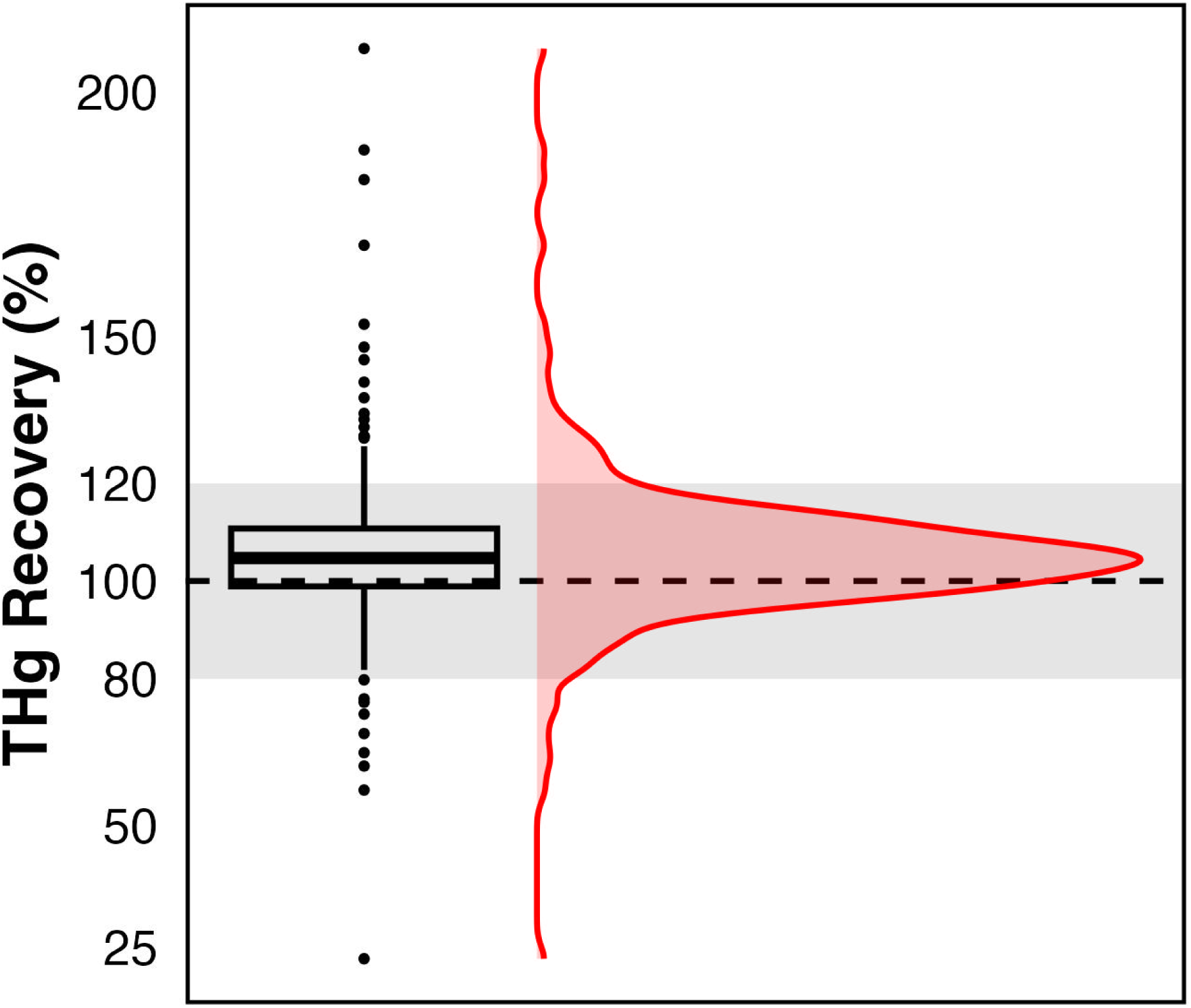
Boxplot and histogram of total mercury (THg) recovery, or the ratio between DBS and whole blood THg concentrations. The boxplot displays the median of the distribution as the bold central line, 25% and 75% quartiles as the box edges, 95% confidence interval as the whisker ends, and outliers as dots. The black dashed line represents a perfect 1:1 relationship between DBS and whole blood THg concentrations, while the gray shaded region represents acceptable accuracy based on United States Environmental Protection Agency Method 7473 (±20% of 100%; USEPA, 1998).

**Figure S6.**
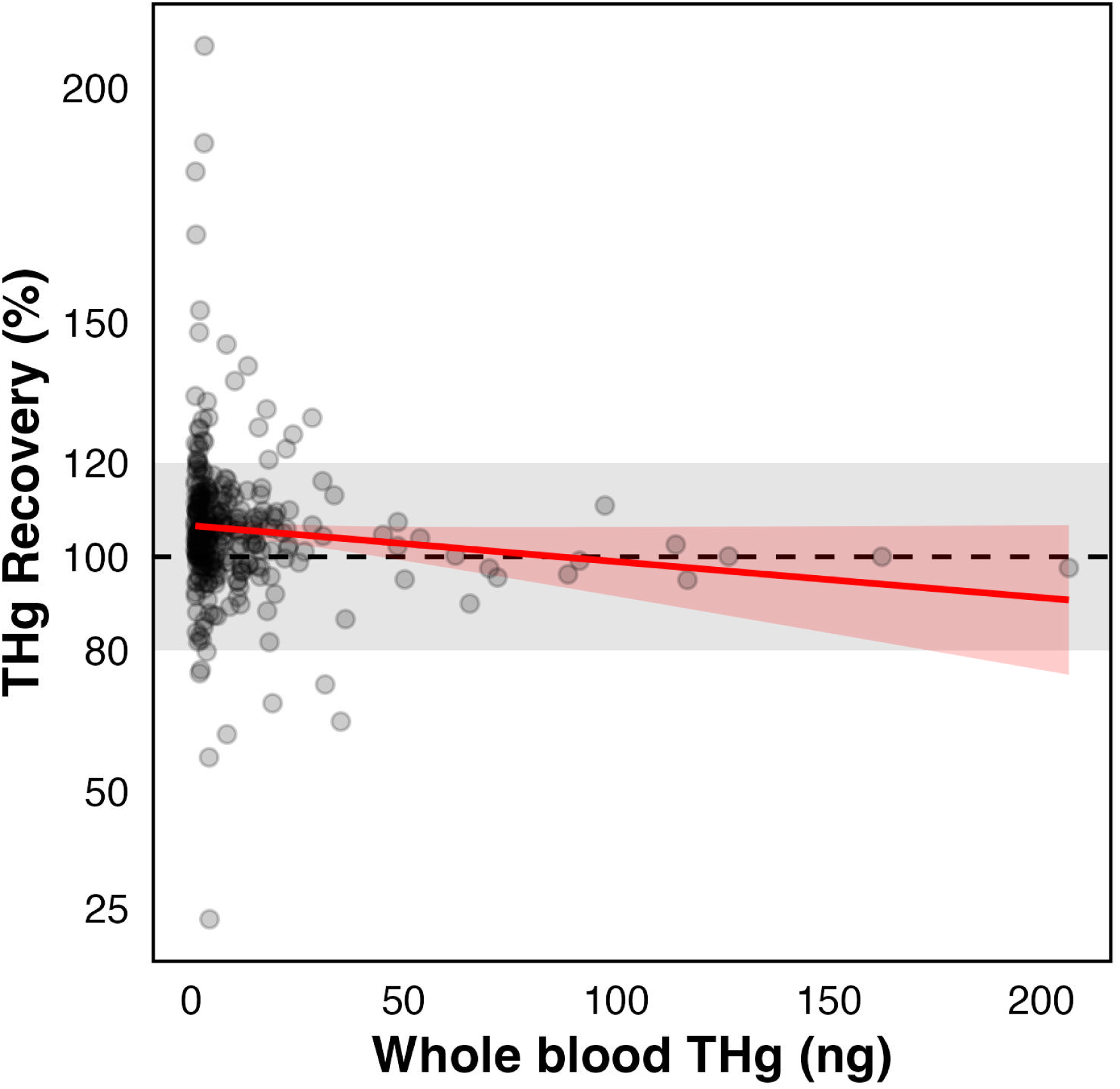
Linear regression of total mercury (THg) recovery and whole blood THg (ng). The black dashed line represents a perfect 1:1 relationship between DBS and whole blood THg concentrations, while the gray shaded region represents acceptable accuracy based on United States Environmental Protection Agency Method 7473 (±20% of 100%; USEPA, 1998).

## References

Ackerman, J. T., Eagles-Smith, C. A., Herzog, M. P., Hartman, C. A., Peterson, S. H., Evers, D. C., Jackson, A. K., Elliott, J. E., Vander Pol, S. S., & Bryan, C. E. (2016). Avian mercury exposure and toxicological risk across western North America: A synthesis. Science of the Total Environment, 568, 749–769. 10.1016/j.scitotenv.2016.03.071

Bang I. (1913). Der Blutzucker. Wiesbaden, Germany.

Barst, B. D., Wooller, M. J., O’Brien, D. M., Santa-Rios, A., Basu, N., Köck, G., Johnson, J. J., & Muir, D. C. G. (2020). Dried Blood Spot Sampling of Landlocked Arctic Char (Salvelinus alpinus) for Estimating Mercury Exposure and Stable Carbon Isotope Fingerprinting of Essential Amino Acids. Environmental Toxicology and Chemistry, 39(4), 893–903. 10.1002/etc.4686

Basu, N., Eng, J. W. L., Perkins, M., Santa-Rios, A., Martincevic, G., Carlson, K., & Neitzel, R. L. (2017). Development and application of a novel method to characterize methylmercury exposure in newborns using dried blood spots. Environmental Research, 159, 276–282. 10.1016/j.envres.2017.08.021

Berlin, M. (1963). Renal Uptake, Excretion, and Retention of Mercury. II. A Study in the Rabbit During Infusion of Methyl- and Phenylmercuric Compounds. Archives of Environmental Health: An International Journal, 6(5), 628–633. 10.1080/00039896.1963.10663451

Brooks, M. E., Kristensen, K., Van Benthem, K. J., Magnusson, A., Berg, C. W., Nielsen, A., Skaug, H. J., Mächler, M., & Bolker, B. M. (2017). glmmTMB Balances Speed and Flexibility Among Packages for Zero-inflated Generalized Linear Mixed Modeling. The R Journal, 9, 378–400.

Chaudhuri, S. N., Butala, S. J. M., Ball, R. W., Braniff, C. T., & Rocky Mountain Biomonitoring Consortium. (2009). Pilot study for utilization of dried blood spots for screening of lead, mercury and cadmium in newborns. Journal of Exposure Science and Environmental Epidemiology, 19, 298–316. 10.1038/jes.2008.19

Dethier, E. N., Silman, M., Leiva, J. D., Alqahtani, S., Fernandez, L. E., Pauca, P., Çamalan, S., Tomhave, P., Magilligan, F. J., Renshaw, C. E., & Lutz, D. A. (2023). A global rise in alluvial mining increases sediment load in tropical rivers. Nature, 620(7975), 787–793. 10.1038/s41586-023-06309-9

Díez, S. (2009). Human Health Effects of Methylmercury Exposure. In Reviews of Environmental Contamination and Toxicology (Vol. 198, pp. 111–132). Springer New York. 10.1007/978-0-387-09647-6

Edmonds, S. T., Evers, D. C., Cristol, D. A., Mettke-Hofmann, C., Powell, L. L., McGann, A. J., Armiger, J. W., Lane, O. P., Tessler, D. F., Newell, P., Heyden, K., & O’Driscoll, N. J. (2010). Geographic and seasonal variation in mercury exposure of the declining Rusty Blackbird. Condor, 112(4), 789–799. 10.1525/cond.2010.100145

Evers, D. (2018). The Effects of Methylmercury on Wildlife: A Comprehensive Review and Approach for Interpretation. In D. A. DellaSala & M. I. Goldstein (Eds.), Encyclopedia of the Anthropocene (Vol. 5). Elsevier Inc. 10.1016/B978-0-12-809665-9.09985-7

Evers, D. C., Ackerman, J. T., Åkerblom, S., Bally, D., Basu, N., Bishop, K., Bodin, N., Braaten, H. F. V., Burton, M. E. H., Bustamante, P., Chen, C., Chételat, J., Christian, L., Dietz, R., Drevnick, P., Eagles-Smith, C., Fernandez, L. E., Hammerschlag, N., Harmelin-Vivien, M., … Wu, P. (2024). Global mercury concentrations in biota: their use as a basis for a global biomonitoring framework. Ecotoxicology 2024 33:4, 33(4), 325–396. 10.1007/S10646-024-02747-X

Evers, D. C., Savoy, L. J., DeSorbo, C. R., Regan, K., Persico, C. P., & Sayers II, C. J. (2021). Bird field sampling methods: collection of tissues for mercury analysis. https://briwildlife.org/wp-content/uploads/2021/07/Bird-Field-Sampling-Methods.pdf

Fair, J. M., Paul, E., Jones, J., Barrett Clark, A., Davie, C., & Kaiser, G. (2010). The Use of Wild Birds in Research. In J. M. Fair, E. Paul, & J. Jones (Eds.), The Ornithological Council (Third edit). 10.2307/1369129

Fang, X., Li, B., Alkhatib, H., Zeng, W., & Yao, Y. (2017). Bayesian inference for the Errors-In-Variables model. Studia Geophysica et Geodaetica, 61(1), 35–52. 10.1007/s11200-015-6107-9

Funk, W. E., McGee, J. K., Olshan, A. F., & Ghio, A. J. (2013). Quantification of arsenic, lead, mercury and cadmium in newborn dried blood spots. Biomarkers, 18(2), 174–177. 10.3109/1354750X.2012.750379

Funk, W. E., Pleil, J. D., Sauter, D. J., McDade, T. W., & Holl, J. L. (2015). Use of Dried Blood Spots for Estimating Children’s Exposures to Heavy Metals in Epidemiological Research. Journal of Environmental & Analytical Toxicology, S7. 10.4172/2161-0525.s7-002

González-Rubio, J. M., Domínguez-Morueco, N., Pedraza-Díaz, S., Cañas Portilla, A., Lucena, M. Á., Rodriguez, A., Castaño, A., & Esteban-López, M. (2023). A simple method for direct mercury analysis in dried blood spots (DBS) samples for human biomonitoring studies. Environment International, 177(December 2022), 107958 Contents. 10.1016/j.envint.2023.107958

Gworek, B., Dmuchowski, W., & Baczewska-Dąbrowska, A. H. (2020). Mercury in the terrestrial environment: a review. Environmental Sciences Europe, 32(1), 128. 10.1186/s12302-020-00401-x

Henderson, L. O., Powell, M. K., Hannon, W. H., Bernert, J. T., Pass, K. A., Fernhoff, P., Ferre, C. D., Martin, L., Franko, E., Rochat, R. W., Brantley, M. D., & Sampson, E. (1997). An Evaluation of the Use of Dried Blood Spots from Newborn Screening for Monitoring the Prevalence of Cocaine Use among Childbearing Women. Biochemical and Molecular Medicine, 61, 143–151.

Holdridge, L. R. (1967). Life Zone Ecology.

Horvat, M., & Byrne, A. R. (1992). Preliminary study of the effects of some physical parameters on the stability of methylmercury in biological samples. The Analyst, 117, 665–668. 10.1039/an9921700665

Jackson, A. K., Evers, D. C., Etterson, M. A., Condon, A. M., Folsom, S. B., Detweiler, J., Schmerfeld, J., & Cristol, D. A. (2011). Mercury exposure affects the reproductive success of a free-living terrestrial songbird, the Carolina wren (Thryothorus ludovicianus). The Auk, 128(4), 759–769. 10.1525/auk.2011.11106

Kershaw, T. G., Dhahir, P. H., & Clarkson, T. W. (1980). The relationship between blood levels and dose of methylmercury in man. Archives of Environmental Health, 35(1), 28–36. 10.1080/00039896.1980.10667458

Langer, E. K., Johnson, K. J., Shafer, M. M., Gorski, P., Overdier, J., Musselman, J., & Ross, J. A. (2010). Characterization of the elemental composition of newborn blood spots using sector-field inductively coupled plasma-mass spectrometry. Journal of Exposure Science & Environmental Epidemiology, 21(4), 355. 10.1038/JES.2010.19

Lehner, A. F., Rumbeiha, W., Shlosberg, A., Stuart, K., Johnson, M., Domenech, R., & Langner, H. (2013). Diagnostic Analysis of Veterinary Dried Blood Spots for Toxic Heavy Metals Exposure. Journal of Analytical Toxicology, 37, 406–422. 10.1093/jat/bkt048

McGuill, M. W., & Rowan, A. N. (1989). Biological Effects of Blood Loss: Implications for Sampling Volumes and Techniques. ILAR Journal, 31(4), 5–20. 10.1093/ilar.31.4.5

Mueller, L. K., Ågerstrand, M., Backhaus, T., Diamond, M., Erdelen, W. R., Evers, D., Groh, K. J., Scheringer, M., Sigmund, G., Wang, Z., & Schäffer, A. (2022). Policy options to account for multiple chemical pollutants threatening biodiversity. Environmental Science: Advances, 2, 151–161. 10.1039/d2va00257d

Norseth, T., & Clarkson, T. W. (1970). Studies on the Biotransformation of 203Hg-labeled Methyl Mercury Chloride in Rats. Archives of Environmental Health, 21(6), 717–727. 10.1080/00039896.1970.10667325

Nyanza, E. C., Dewey, D., Bernier, F., Manyama, M., Hatfield, J., & Martin, J. W. (2019). Validation of dried blood spots for maternal biomonitoring of nonessential elements in an artisanal and small-scale gold mining area of Tanzania. Environmental Toxicology and Chemistry, 38(6), 1285–1293. 10.1002/etc.4420

Paul, E. (2005). A guide to the permits and procedures for importing bird products into the United States for scientific research and display. The Ornithological Council, 97. http://naturalhistory.si.edu/BIRDNET/documents/importguidespecimens.pdf

Perkins, M., & Basu, N. (2018). Dried blood spots for estimating mercury exposure in birds. Environmental Pollution, 236, 236–246. 10.1016/j.envpol.2018.01.036

Reilly, P. M., & Patino-Lea, H. (2012). A Bayesian Study of the Error-in-Variables Model. Technometrics, 23, 221–231. 10.1080/00401706.1981.10486290

Rimmer, C. C., Mcfarland, K. P., Evers, D. C., Miller, E. K., Aubry, Y., Busby, D., & Taylor, R. J. (2005). Mercury concentrations in Bicknell’s thrush and other insectivorous passerines in montane forests of northeastern North America. Ecotoxicology, 14, 223–240. papers3://publication/uuid/B9A71908-3061-4884-BB83-38F08E5BCC5C

Santa-Rios, A., Barst, B. D., Tejeda-Benitez, L., Palacios-Torres, Y., Baumgartner, J., & Basu, N. (2021). Dried blood spots to characterize mercury speciation and exposure in a Colombian artisanal and small-scale gold mining community. Chemosphere, 266, 129001. 10.1016/j.chemosphere.2020.129001

Sayers II, C. J., Evers, D. C., Ruiz-Gutierrez, V., Adams, E., Vega, C. M., Pisconte, J. N., Tejeda, V., Regan, K., Lane, O. P., Ash, A. A., Cal, R., Reneau, S., Martínez, W., Welch, G., Hartwell, K., Teul, M., Tzul, D., Arendt, W. J., Tórrez, M. A., … Gerson, J. (2023). Mercury in Neotropical birds: a synthesis and prospectus on 13 years of exposure data. Ecotoxicology, 32(8), 1096–1123. 10.1007/s10646-023-02706-y

Sayers II, C. J., Roeder, M. R., Forrette, L. M., Roche, D., Dupont, G. L. B., Apgar, S. E., Kocek, A. R., Cook, A. M., Shriver, W. G., Elphick, C. S., Olsen, B., & Bonter, D. N. (2021). Geographic variation of mercury in breeding tidal marsh sparrows of the northeastern United States. Ecotoxicology, 30(9), 1929–1940. 10.1007/s10646-021-02461-y

Shlosberg, A., Rumbeiha, W. K., Lublin, A., & Kannan, K. (2011). A database of avian blood spot examinations for exposure of wild birds to environmental toxicants: The DABSE biomonitoring project. Journal of Environmental Monitoring, 13(6), 1547–1558. 10.1039/c0em00754d

Shlosberg, A., Wu, Q., Rumbeiha, W. K., Lehner, A., Cuneah, O., King, R., Hatzofe, O., Kannan, K., & Johnson, M. (2012a). Examination of Eurasian Griffon Vultures (Gyps fulvus fulvus) in Israel for Exposure to Environmental Toxicants Using Dried Blood Spots. Archives of Environmental Contamination and Toxicology, 62(3), 502–511. 10.1007/s00244-011-9709-4

Shlosberg, A., Wu, Q., Rumbeiha, W. K., Lehner, A., Cuneah, O., King, R., Hatzofe, O., Kannan, K., & Johnson, M. (2012b). Examination of Eurasian griffon vultures (Gyps fulvus fulvus) in Israel for exposure to environmental toxicants using dried blood spots. Archives of Environmental Contamination and Toxicology, 62, 502–511. 10.1007/s00244-011-9709-4

Shoari, N., & Dubé, J. S. (2018). Toward Improved Analysis of Concentration Data: Embracing Nondetects. Environmental Toxicology and Chemistry, 37, 643–656. 10.1002/etc.4046

Sjöholm, M. I. L., Dillner, J., & Carlson, J. (2007). Assessing quality and functionality of DNA from fresh and archival dried blood spots and recommendations for quality control guidelines. Clinical Chemistry, 53(8), 1401–1407. 10.1373/clinchem.2007.087510

Sommer, Y. L., Ward, C. D., Pan, Y., Caldwell, K. L., & Jones, R. L. (2016). Long-term stability of inorganic, methyl and ethyl mercury in whole blood: effects of storage temperature and time. Journal of Analytical Toxicology, 40, 222–228. 10.1093/jat/bkw007

Stotz, D. F., Fitzpatrick, J. W., Parker III, T. A., & Moskovits, D. K. (1996). Neotropical Birds: Ecology and Conservation. University of Chicago Press.

Stove, C. P., Ingels, A. S. M. E., de Kesel, P. M. M., & Lambert, W. E. (2012). Dried blood spots in toxicology: From the cradle to the grave? Critical Reviews in Toxicology, 42(3), 230–243. 10.3109/10408444.2011.650790

UN (United Nations). (2013). Minamata Convention on Mercury. https://treaties.un.org/doc/Treaties/2013/10/20131010%2011-16%20AM/CTC-XXVII-17.pdf

USEPA (United States Environmental Protection Agency). (1998). Method 7473 (SW-846): Mercury in Solids and Solutions by Thermal Decompostion, Amalgamation, and Atomic Absorption Spectrophotometry. https://www.epa.gov/sites/default/files/2015-07/documents/epa-7473.pdf

Varian-Ramos, C. W., Condon, A. M., Hallinger, K. K., Carlson-Drexler, K. A., & Cristol, D. A. (2011). Stability of mercury concentrations in frozen avian blood samples. Bulletin of Environmental Contamination and Toxicology, 86, 159–162. 10.1007/s00128-010-0164-0

WGC (World Gold Council). (2026). Gold Demand Trends: Q4 and Full Year 2025. https://www.gold.org/goldhub/data/gold-demand-by-country

Whitney, M. C., & Cristol, D. A. (2017). Impacts of sublethal mercury exposure on birds: a detailed review. In P. de Voogt (Ed.), Reviews of Environmental Contamination and Toxicology (Reviews of, Vol. 244, pp. 113–163). Springer Nature.

