## Supplementary material for "A field method to optimize dried blood spot sampling for mercury biomonitoring": code.R

```
# Authors:
# Last updated: 24 May 2025

# Full analysis and supplemental materials for 
# "A field method to optimize dried blood spot sampling for mercury biomonitoring"

# Contents ----------------------------------------------------------------

# Load packages & data
# Part 1: Summary statistics
# Part 2: Bayesian linear regression
# Part 2a: JAGS model
# Part 2b: Model summary
# Part 2c: Posterior predictive check
# Part 2d: Prediction
# Part 3: Hg Recovery

# Load packages & data ----------------------------------------------------

library(tidyverse)
library(readxl)
library(janitor)
library(ggpubr)
library(ggview)
library(R2jags)
library(foreign)
library(dclone)
library(MCMCvis)
library(lattice)
library(glmmTMB)
library(beepr)

# Resolve namespace conflicts and create necessary functions
select <- dplyr::select
"%nin%" <- Negate("%in%")

# Import THg data
THg <- read_csv("Data/THg.csv") %>% 
  # Remove samples below the limits of detection of our instruments
  mutate(THg_ng_DBS = if_else(Analyzer == "Milestone DMA-80" & THg_ng_DBS < 0.01, NA, THg_ng_DBS),
         THg_ng_Blood = if_else(Analyzer == "Milestone DMA-80" & THg_ng_Blood < 0.01, NA, THg_ng_Blood),
         THg_ng_DBS = if_else(Analyzer == "Nippon MA-3000" & THg_ng_DBS < 0.001, NA, THg_ng_DBS),
         THg_ng_Blood = if_else(Analyzer == "Nippon MA-3000" & THg_ng_Blood < 0.001, NA, THg_ng_Blood)) %>% 
  filter(!is.na(THg_ng_DBS) & !is.na(THg_ng_Blood))

# Import blank data
blanks <- read_csv("Data/blanks.csv") %>% 
  # Remove samples below the limits of detection of our instruments
  mutate(THg_ng = if_else(Lab == "LAMQA" & THg_ng < 0.01, NA, THg_ng),
         THg_ng = if_else(Lab == "BRI Toxicology Lab" & THg_ng < 0.001, NA, THg_ng)) %>% 
  filter(!is.na(THg_ng))


# Part 1: Summary statistics ----------------------------------------------

# Sample size
nrow(THg) # Number of individual samples
length(unique(THg$Species)) # Number of species
length(unique(THg$Site)) # Number of sites
hist(THg$Volume_µL_DBS)

# Calculate percent of samples from each laboratory
THg %>% 
  group_by(Lab) %>% 
  summarize(n = n())

# Simple linear regression of raw data
cor(THg$THg_ppm_DBS, THg$THg_ppm_Blood, method = "pearson")
cor(THg$log_THg_ppm_DBS, THg$log_THg_ppm_Blood, method = "pearson")

ggplot() + 
  geom_ribbon(aes(x = seq(0, 10), ymin = seq(0, 10) * 0.8, ymax = seq(0, 10) * 1.2),
              fill = "gray", alpha = 0.4) +
  geom_abline(intercept = 0, slope = 1.0, linetype = "dashed") +
  geom_point(data = THg, mapping = aes(x = THg_ppm_Blood, y = THg_ppm_DBS), alpha = 0.2, size = 2) +
  geom_smooth(data = THg, mapping = aes(x = THg_ppm_Blood, y = THg_ppm_DBS),
              method = "lm", color = "red", fill = "red", se = T, level = 0.95, linewidth = 1) +
  coord_cartesian(xlim = c(0, 8), ylim = c(0, 8)) +
  theme_minimal(base_size = 16) +
  labs(y = "Dried blood spot THg (µg/L)", x = "Whole blood THg (µg/g ww)") +
  theme(axis.title.x = element_text(face = "bold"),
        axis.title.y = element_text(face = "bold"),
        axis.text = element_text(color = "black"),
        panel.border = element_rect(color = "black", fill = NA, linewidth = 1),
        aspect.ratio = 1) +
  canvas(units = "in", width = 5, height = 5, dpi = 300)

# Blanks
blanks %>% 
  group_by(Lab) %>% 
  summarize(n = n(),
            Mean = mean(THg_ppm),
            SD = sd(THg_ppm),
            Min = min(THg_ppm),
            Max = max(THg_ppm))


# Part 2: Bayesian linear regression --------------------------------------
# Part 2a: JAGS model -----------------------------------------------------

jags_data <- list(obs_DBS = THg$THg_ppm_DBS,
                  obs_WB = THg$THg_ppm_Blood,
                  Site = as.factor(THg$Site),
                  Species = as.factor(THg$Species),
                  Analyzer = as.numeric(as.factor(THg$Analyzer)), # 1 = Milestone DMA-80, 2 = Nippon MA-3000
                  n_sample = length(THg$THg_ppm_DBS),
                  n_site = length(unique(THg$Site)),
                  n_spp = length(unique(THg$Species)),
                  n_analyzer = length(unique(THg$Analyzer)))

lm1 <- function() {
  
  # Priors
  intercept <- 0
  slope ~ dnorm(0, 0.0001)
  
  tau_site ~ dgamma(0.0001, 0.0001)
  tau_spp ~ dgamma(0.0001, 0.0001)
  
  sigma_site <- 1 / sqrt(tau_site)
  sigma_spp <- 1 / sqrt(tau_spp)
  
  # Random slopes
  for (j in 1:n_site) {
    site[j] ~ dnorm(0, tau_site)
  }
  
  for (k in 1:n_spp) {
    spp[k] ~ dnorm(0, tau_spp)
  }
  
  # Instrument-specific measurement error priors
  for (a in 1:n_analyzer) {
    tau_analyzer[a] ~ dgamma(0.0001, 0.0001) # One precision parameter per instrument
    sigma_analyzer[a] <- 1 / sqrt(tau_analyzer[a])
  }
  
  # Likelihood
  for (i in 1:n_sample) {
    
    # Stochastic processes
    # Measurement error is indexed by which instrument analyzed sample i
    # Observed DBS concentrations measured with error around their latent truth
    obs_DBS[i] ~ dnorm(true_DBS[i], tau_analyzer[Analyzer[i]])
    
    # Latent whole blood concentration mean
    true_WB[i] ~ dnorm(obs_WB[i], tau_analyzer[Analyzer[i]])
    #obs_WB[i] ~ dnorm(true_WB[i], tau_analyzer[Analyzer[i]])
    
    # Deterministic process for latent DBS mean
    true_DBS[i] <- intercept + # Intercept fixed at 0
      (slope + site[Site[i]] + spp[Species[i]]) * true_WB[i] # Slope + random slopes for species and site
  }
}

lm1_fit <- jags(data = jags_data,
                parameters.to.save = c("true_DBS",
                                       "true_WB",
                                       "slope",
                                       "sigma_analyzer",
                                       "sigma_site",
                                       "site",
                                       "sigma_spp",
                                       "spp"),
                model.file = lm1,
                n.chains = 3,
                n.iter = 55000,
                n.burnin = 5000,
                n.thin = 50)

save(lm1_fit, file = "Data/DBS_R2Jags.RData")
load("Data/DBS_R2Jags.RData")


# Part 2b: Model summary -------------------------------------------------

# Summary of posterior distributions
# R-hats are all below 1.1 with good effective n
# Variance of Milestone DMA-80 is double that of Nippon MA-3000, but both are small
MCMCsummary(lm1_fit, params = c("slope",
                                "sigma_analyzer",
                                "sigma_site",
                                "sigma_spp"
),
round = 2)

# Trace plots of MCMC chains
MCMCtrace(lm1_fit, params = c("slope",
                              "sigma_analyzer",
                              "sigma_site",
                              "sigma_spp"
),
priors = rnorm(27000, 0, sqrt(1/0.001)),
Rhat = TRUE, n.eff = TRUE, pdf = FALSE)

# Effect plot for fixed effects
MCMCplot(lm1_fit,
         params = c("slope",
                    "sigma_analyzer",
                    "sigma_site",
                    "sigma_spp"
         ),
         labels = c("Slope: Whole blood THg",
                    "Variance: Milestone DMA-80",
                    "Variance: Nippon MA-3000",
                    "Variance: Site",
                    "Variance: Species"
         ),
         #ci = c(50, 95),
         ref_ovl = TRUE)

# Effect plot for site
jpeg(filename = "Figures/FigS2_bayesian_site.jpg", units = "in", width = 6, height = 4, res = 300, quality = 100)
MCMCplot(lm1_fit,
         params = "site",
         labels = unique(THg$Site),
         rank = TRUE,
         ci = c(0.1, 95),
         ref_ovl = TRUE)
dev.off()

# Effect plot for species
jpeg(filename = "Figures/FigS3_bayesian_spp.jpg", units = "in", width = 8, height = 10, res = 300, quality = 100)
MCMCplot(lm1_fit,
         params = "spp",
         labels = unique(THg$Species),
         rank = TRUE,
         ci = c(0.01, 95),
         ref_ovl = TRUE)
dev.off()


# Part 2c: Posterior predictive check -------------------------------------

# Extract all relevant parameters
mcmc <- as.matrix(lm1_fit$BUGSoutput$sims.matrix)
slope <- mcmc[, "slope"]
true_WB_mat <- mcmc[, grep("^true_WB\\[", colnames(mcmc))] # Extract full true_WB matrix (n_draws x n_obs)
sigma_analyzer <- mcmc[, grep("^sigma_analyzer\\[", colnames(mcmc))]

# Extract site and species random effects
site_effects <- mcmc[, grep("^site\\[", colnames(mcmc))]
spp_effects <- mcmc[, grep("^spp\\[", colnames(mcmc))]

# Integer indices for grouping variables
site_idx <- as.numeric(as.factor(THg$Site))
spp_idx <- as.numeric(as.factor(THg$Species))
analyzer_idx <- as.numeric(as.factor(THg$Analyzer))

# Reconstruct true_DBS for each posterior draw
n_draws <- nrow(mcmc)
n_obs <- nrow(THg)
true_DBS_mat <- matrix(NA, nrow = n_draws, ncol = n_obs)

for (i in 1:n_draws) {
  true_DBS_mat[i, ] <- 
    (slope[i] + site_effects[i, site_idx] + spp_effects[i, spp_idx]) * true_WB_mat[i, ]
  
}

# Draw replicated data from the posterior predictive distribution
y_rep <- matrix(NA, nrow = n_draws, ncol = n_obs)
for (i in 1:n_draws) {
  y_rep[i, ] <- rnorm(n_obs, true_DBS_mat[i, ], sigma_analyzer[i, analyzer_idx])
}

# Subsample to 500 draws to keep plotting manageable
draw_x <- sample(1:n_draws, 500)
y_rep_sub <- y_rep[draw_x, ]

# Plot
ggplot() +
  geom_density(aes(x = as.vector(y_rep_sub), fill = "Predicted"), alpha = 0.5) +
  geom_density(aes(x = THg$THg_ppm_Blood, fill = "Observed"), alpha = 0.5) +
  theme_minimal(base_size = 16) +
  #labs(x = expression(THg[DBS]), y = "Density") +
  labs(x = "Dried blood spot THg (µg/L)", y = "Density") +
  theme(axis.title.x = element_text(face = "bold"),
        axis.title.y = element_text(face = "bold"),
        axis.text = element_text(color = "black"),
        legend.title = element_blank(),
        legend.position = "right",
        panel.border = element_rect(color = "black", fill = NA, linewidth = 1),
        aspect.ratio = 1) +
  canvas(units = "in", width = 6, height = 4.5, dpi = 300)

ggsave("Figures/FigS1_PPC.jpg", dpi = 300, width = 6, height = 4.5)


# R-squared
# Variance of predicted values for each draw
var_pred <- apply(true_DBS_mat, 1, var)

# Variance of residuals for each draw
resid_mat <- matrix(NA, nrow = n_draws, ncol = n_obs)
for (i in 1:n_draws) {
  resid_mat[i, ] <- THg$THg_ppm_DBS - true_DBS_mat[i, ]
}
var_resid <- apply(resid_mat, 1, var)

# Bayesian R-squared for each draw
r2_bayes <- var_pred / (var_pred + var_resid)

# Summarize
mean(r2_bayes)
median(r2_bayes)
quantile(r2_bayes, c(0.025, 0.5, 0.975))
hist(r2_bayes, breaks = 50, main = "Bayesian R²", xlab = "R²")


# Part 2d: Prediction -----------------------------------------------------

# Prediction sequence on raw scale, evenly spaced on log scale
n_pred <- 1000
x_seq <- seq(min(THg$THg_ppm_Blood), max(THg$THg_ppm_Blood), length.out = n_pred)

# Extract posterior samples
slope <- mcmc[, "slope"]
n_draws <- length(slope)

# Build prediction matrix (n_draws x n_pred)
# Each row is one posterior draw, each column is one x value
# Marginal prediction: ignore random effects
pred_mat <- outer(slope, x_seq)

# Summarize across draws
mean_pred <- apply(pred_mat, 2, mean)
ci_low <- apply(pred_mat, 2, quantile, probs = 0.025)
ci_high <- apply(pred_mat, 2, quantile, probs = 0.975)
pred_df <- data.frame(x_seq = x_seq,
                      y_hat = mean_pred,
                      ci_low = ci_low,
                      ci_high = ci_high)

# Plot
ggplot() +
  geom_ribbon(aes(x = seq(0, 10), ymin = seq(0, 10) * 0.8, ymax = seq(0, 10) * 1.2), 
              fill = "gray", alpha = 0.4) +
  geom_abline(intercept = 0, slope = 1.0, linetype = "dashed", color = "black") +
  geom_ribbon(data = pred_df, aes(x = x_seq, ymin = ci_low, ymax = ci_high), 
              fill = "red", alpha = 0.2) +
  geom_point(data = THg, aes(x = THg_ppm_Blood, y = THg_ppm_DBS), 
             color = "black", alpha = 0.2, size = 2) +
  geom_line(data = pred_df, aes(x = x_seq, y = y_hat), 
            color = "red") +
  coord_cartesian(xlim = c(0, 8), ylim = c(0, 8)) +
  theme_minimal(base_size = 16) +
  labs(x = "Whole blood THg (µg/g ww)", y = "Dried blood spot THg (µg/L)") +
  theme(axis.title.x = element_text(face = "bold"),
        axis.title.y = element_text(face = "bold"),
        axis.text = element_text(color = "black"),
        panel.border = element_rect(color = "black", fill = NA, linewidth = 1),
        aspect.ratio = 1) +
  canvas(units = "in", width = 5, height = 5, dpi = 300)

ggsave("Figures/Fig2_DBSvWB_Bayesian.jpg", dpi = 300, width = 5, height = 5)


# Part 3: Hg Recovery -----------------------------------------------------

# Assess Hg recovery (DBS THg/whole blood THg)
recovery <- THg %>% 
  filter(THg_ng_Blood >= 1 & THg_ng_DBS >= 1) %>% 
  mutate(THg_Recovery = THg_ppm_DBS/THg_ppm_Blood)

summary(recovery$THg_Recovery)
sd(recovery$THg_Recovery)

# What percentage of the data violate the USEPA's rule of ±20% deviance?
nrow(recovery %>% filter(THg_Recovery > 1.2 | THg_Recovery < 0.8))/nrow(recovery)

# Recovery boxplot with uncorrected DBS
ggplot() + 
  annotate("rect", ymin = 0.8, ymax = 1.2, xmin = -Inf, xmax = Inf, fill = "gray", alpha = 0.4) +
  geom_hline(yintercept = 1, linetype = "dashed") +
  geom_boxplot(data = recovery, aes(x = -1.3, y = THg_Recovery), color = "black", fill = NA,
               width = 2, size = 0.75) +
  geom_density(data = recovery, aes(y = THg_Recovery), color = "red", fill = "red", alpha = 0.2) +
  theme_minimal(base_size = 16) +
  labs(y = "THg Recovery (%)") +
  scale_x_continuous(breaks = c()) + 
  scale_y_continuous(#labels = scales::percent,
    breaks = c(0.25, 0.5, .8, 1, 1.2, 1.5, 2),
    labels = c(25, 50, 80, 100, 120, 150, 200)
  ) +
  theme(axis.title.x = element_blank(),
        axis.title.y = element_text(face = "bold", color = "black"),
        axis.text.y = element_text(color = "black"),
        axis.ticks = element_blank(),
        legend.title = element_blank(),
        legend.position = "right",
        panel.border = element_rect(color = "black", fill = NA, linewidth = 1),
        panel.grid.major = element_blank(),
        panel.grid.minor = element_blank(),
        aspect.ratio = 1) +
  canvas(units = "in", width = 4.5, height = 4)
ggsave("Figures/FigS4_Recovery.jpg", dpi = 300, width = 4.5, height = 4)


# Variation in THg recovery is highest towards the analytical limit of detection,
# indicating non-uniform error (heteroscedasticity)
plot(recovery$THg_Recovery ~ recovery$Volume_µL_DBS)
plot(recovery$THg_Recovery ~ recovery$Mass_mg_Blood)
plot(recovery$THg_Recovery ~ recovery$THg_ng_DBS)
plot(recovery$THg_Recovery ~ recovery$THg_ppm_DBS)
plot(recovery$THg_Recovery ~ recovery$THg_ng_Blood)
plot(recovery$THg_Recovery ~ recovery$THg_ppm_Blood)

rec <- glmmTMB(THg_Recovery ~ THg_ng_Blood + (1 | Species) + (1 | Site),
               data = recovery, family = "gaussian", REML = F)

summary(rec)
confint(rec)
performance::r2(rec)
car::Anova(freq, type = 2)

n_pred = 1000
THg_ng_Blood <- seq(min(recovery$THg_ng_Blood), max(recovery$THg_ng_Blood), length.out = n_pred)
x_df <- data.frame(THg_ng_Blood)
pred <- predict(rec, newdata = x_df, se.fit = TRUE, re.form = NA)
pred_df <- data.frame(THg_ng_Blood = THg_ng_Blood,
                      y_hat = pred$fit, 
                      ci_low = pred$fit - 1.96 * pred$se.fit,
                      ci_high = pred$fit + 1.96 * pred$se.fit)
ggplot() +
  annotate("rect", ymin = 0.8, ymax = 1.2, xmin = -Inf, xmax = Inf, fill = "gray", alpha = 0.4) +
  geom_hline(yintercept = 1, linetype = "dashed") +
  geom_ribbon(data = pred_df, mapping = aes(x = THg_ng_Blood, ymin = ci_low, ymax = ci_high), fill = "red", alpha = 0.2) +
  geom_point(data = recovery, mapping = aes(x = THg_ng_Blood, y = THg_Recovery), color = "black", alpha = 0.2) +
  geom_line(data = pred_df, mapping = aes(x = THg_ng_Blood, y = y_hat), color = "red", linewidth = 1) +
  theme_minimal(base_size = 16) +
  labs(y = "THg Recovery (%)", x = "Whole blood THg (ng)") +
  scale_x_continuous() +
  scale_y_continuous(#labels = scales::percent,
    breaks = c(0.25, 0.5, .8, 1, 1.2, 1.5, 2),
    labels = c(25, 50, 80, 100, 120, 150, 200)
  ) +
  theme(axis.title.x = element_text(face = "bold"),
        axis.title.y = element_text(face = "bold"),
        axis.text = element_text(color = "black"),
        legend.title = element_blank(),
        legend.position = "right",
        panel.border = element_rect(color = "black", fill = NA, linewidth = 1),
        panel.grid.major = element_blank(),
        panel.grid.minor = element_blank(),
        aspect.ratio = 1) +
  canvas(units = "in", width = 5.5, height = 5, dpi = 300)

ggsave("Figures/FigS5_Recovery_Heteroscadasticity.jpg", dpi = 300, width = 5.5, height = 5)
```
