## Supplementary figures and images for "A field method to optimize dried blood spot sampling for mercury biomonitoring"

### Fig1_DBS-Method.pdf

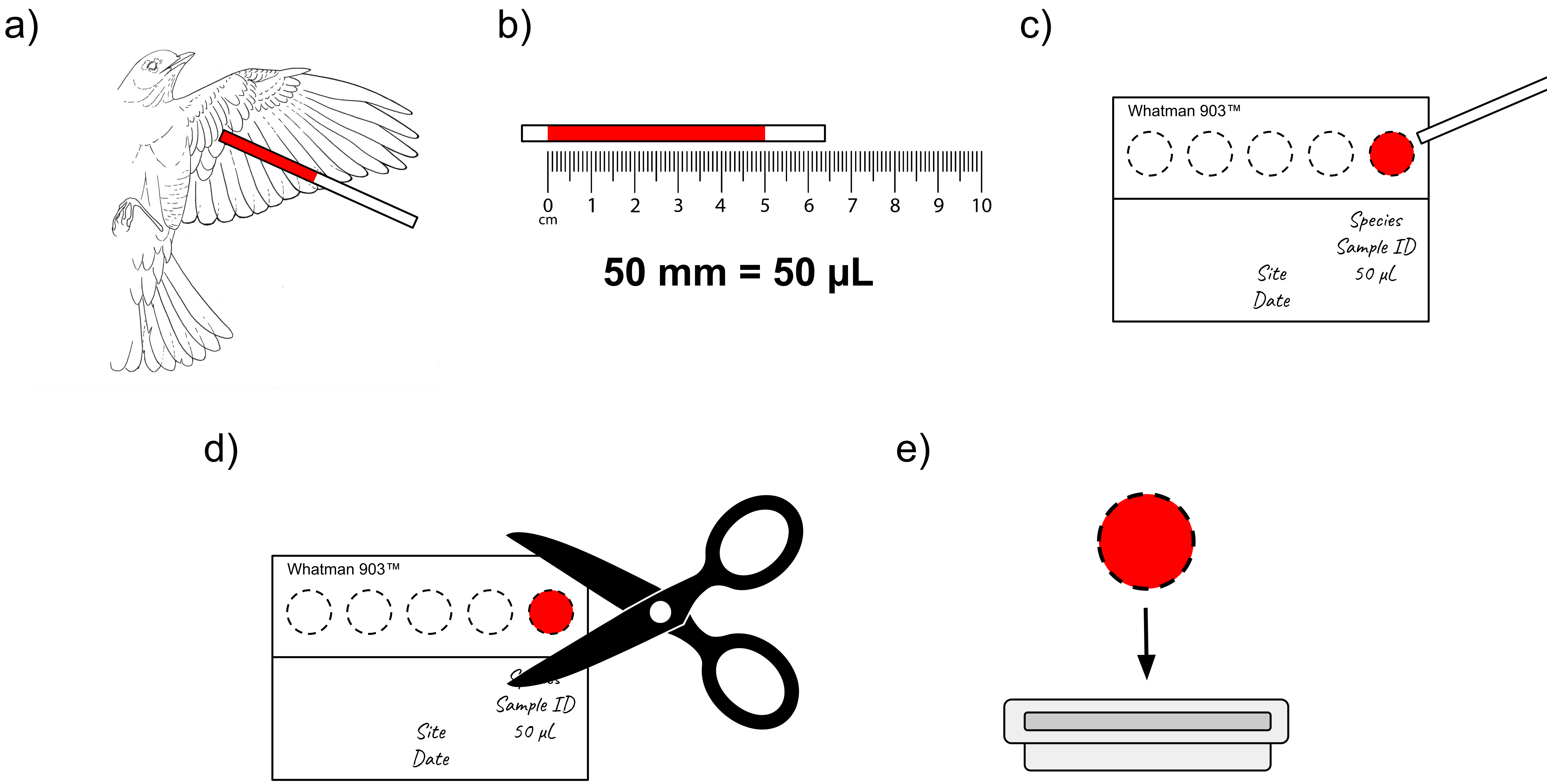

### Fig2_DBSvWB_Bayesian.jpg

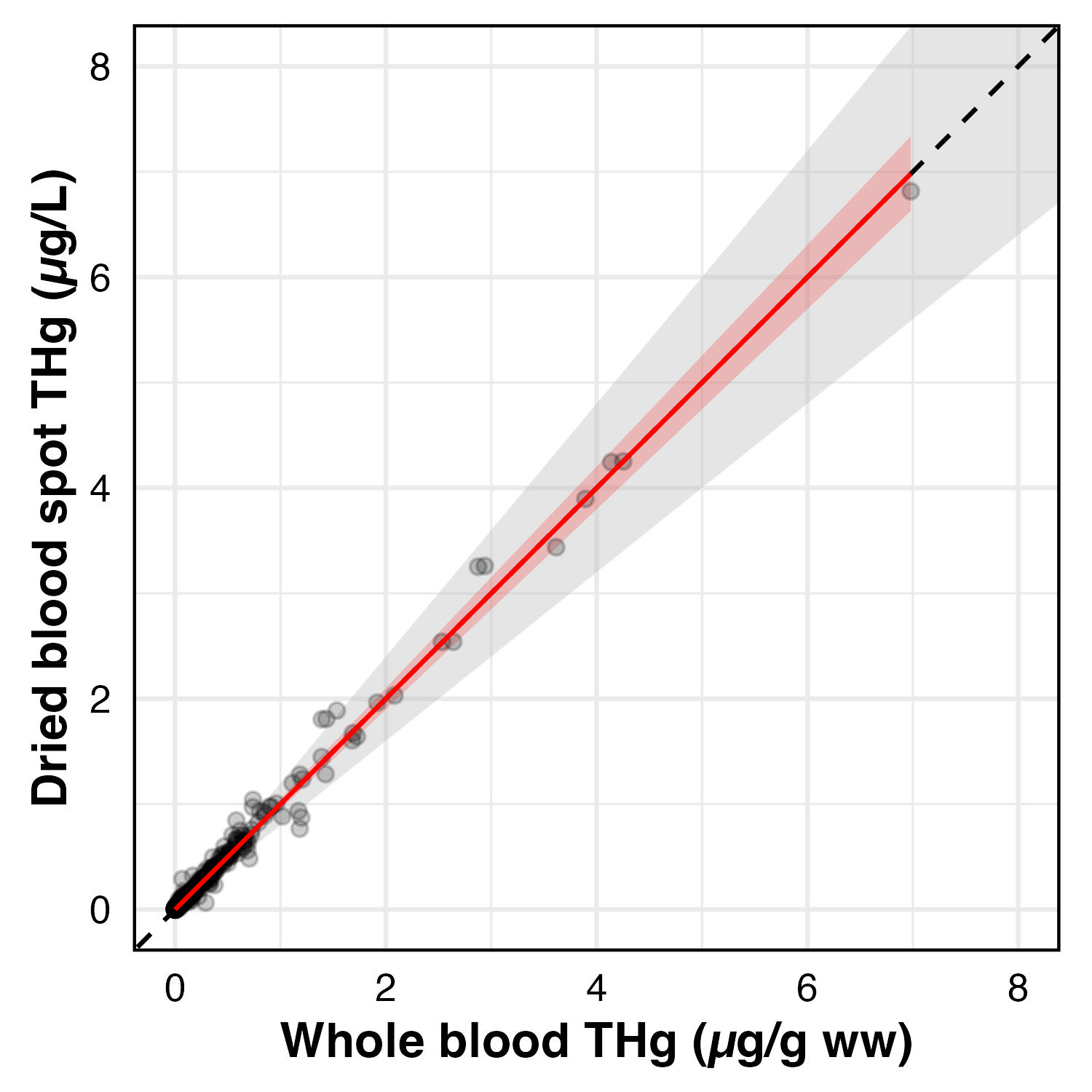

### FigS1_PPC.jpg

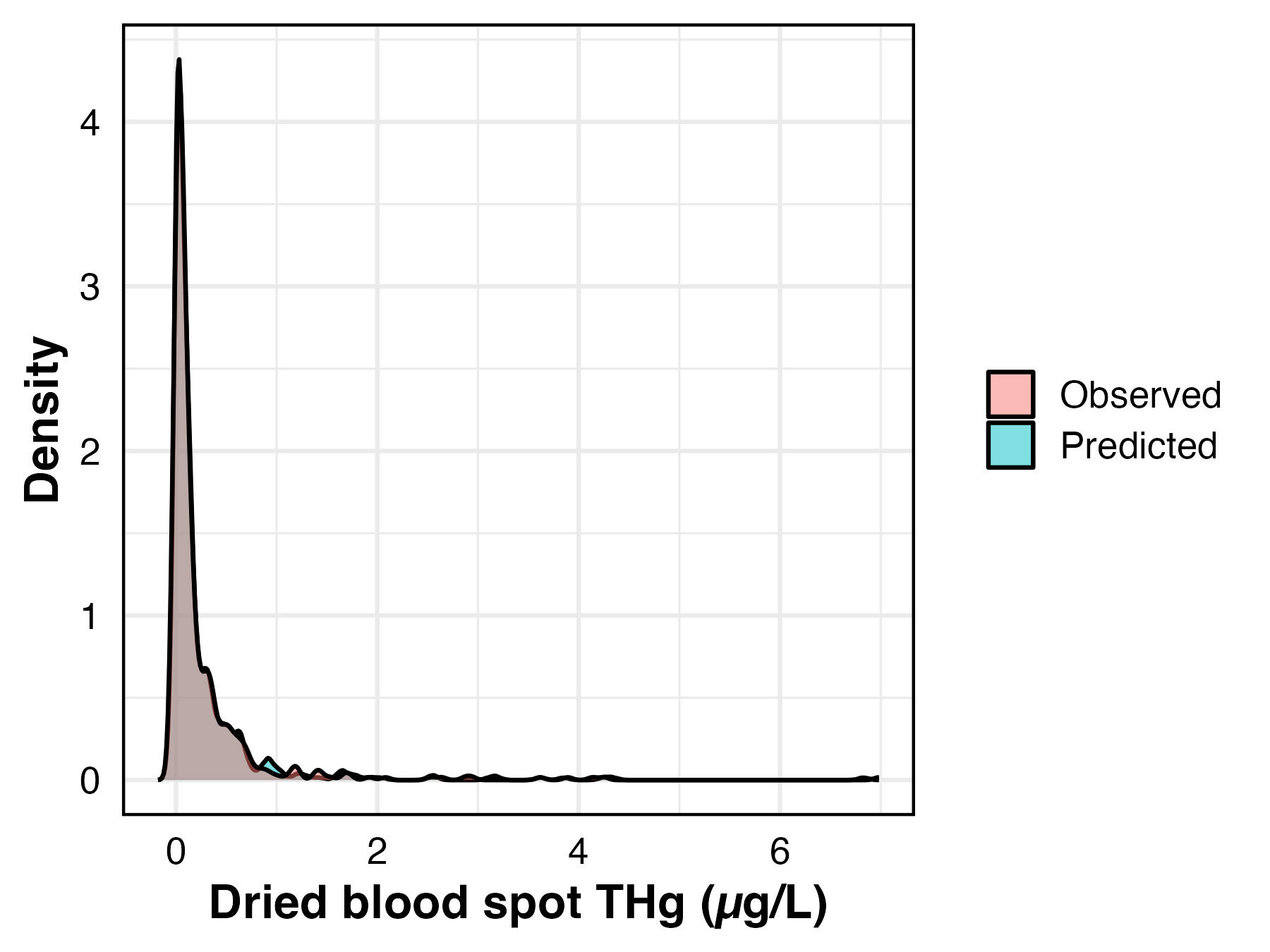

### FigS2_bayesian_site.jpg

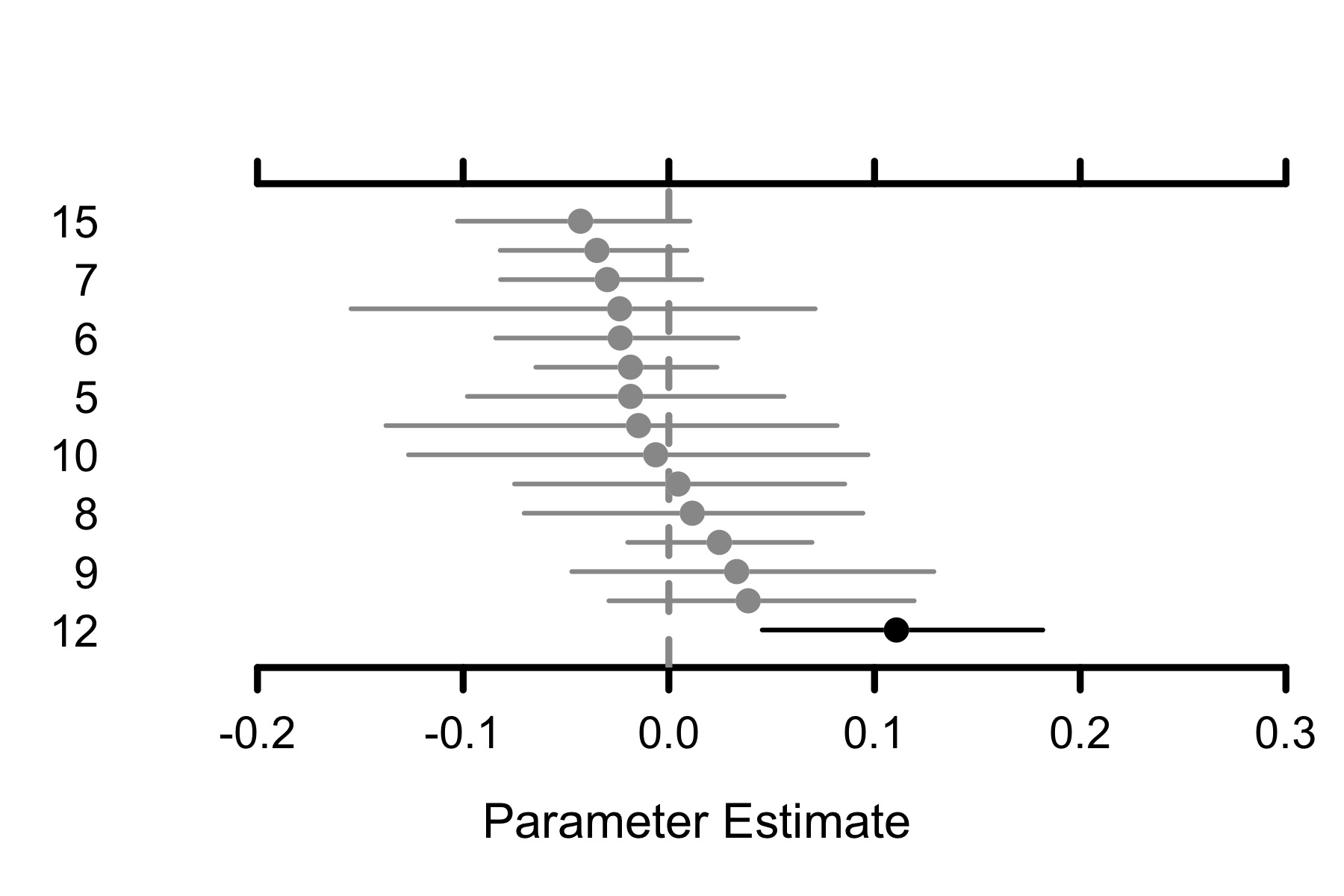

### FigS3_bayesian_spp.jpg

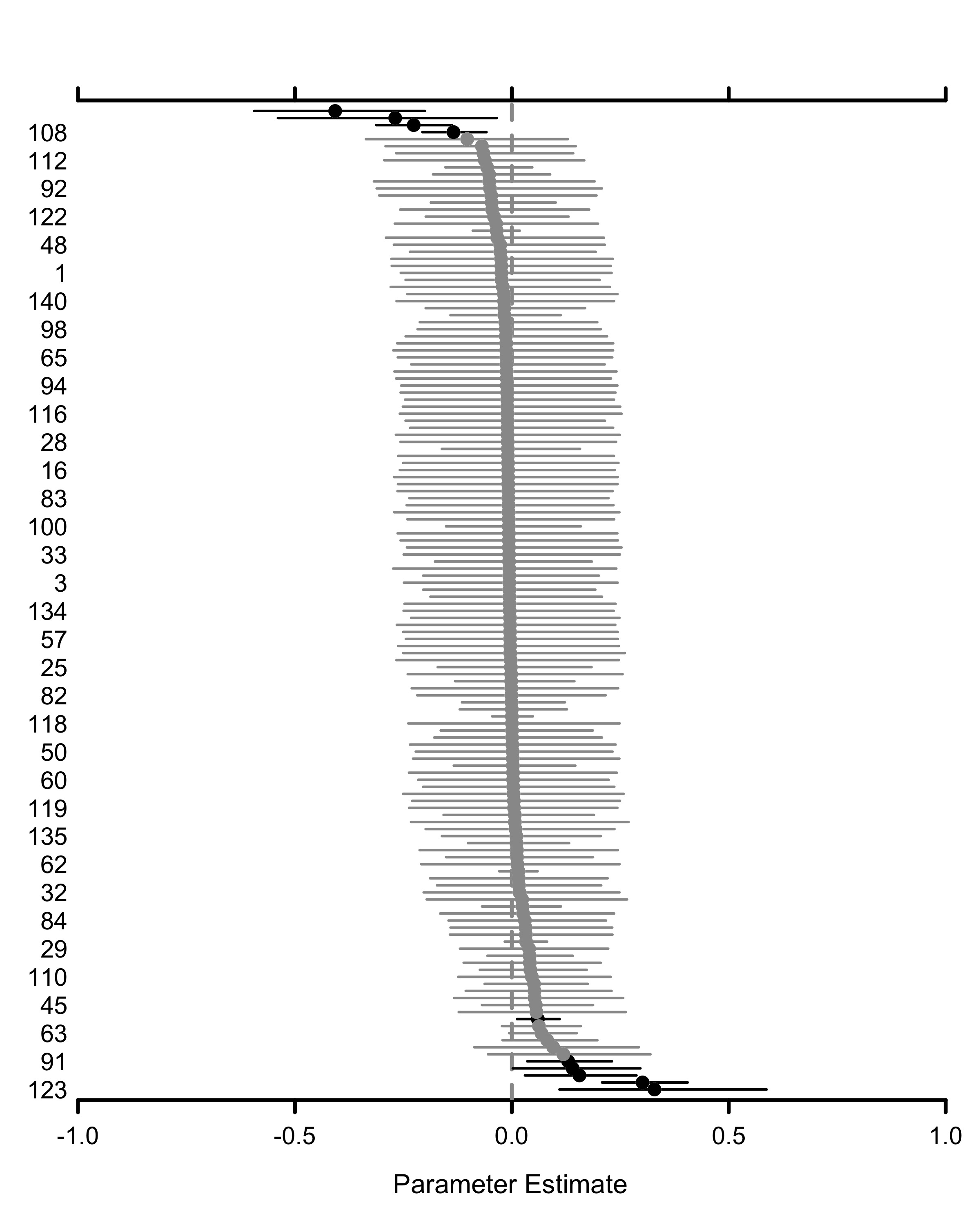

### FigS4_Recovery.jpg

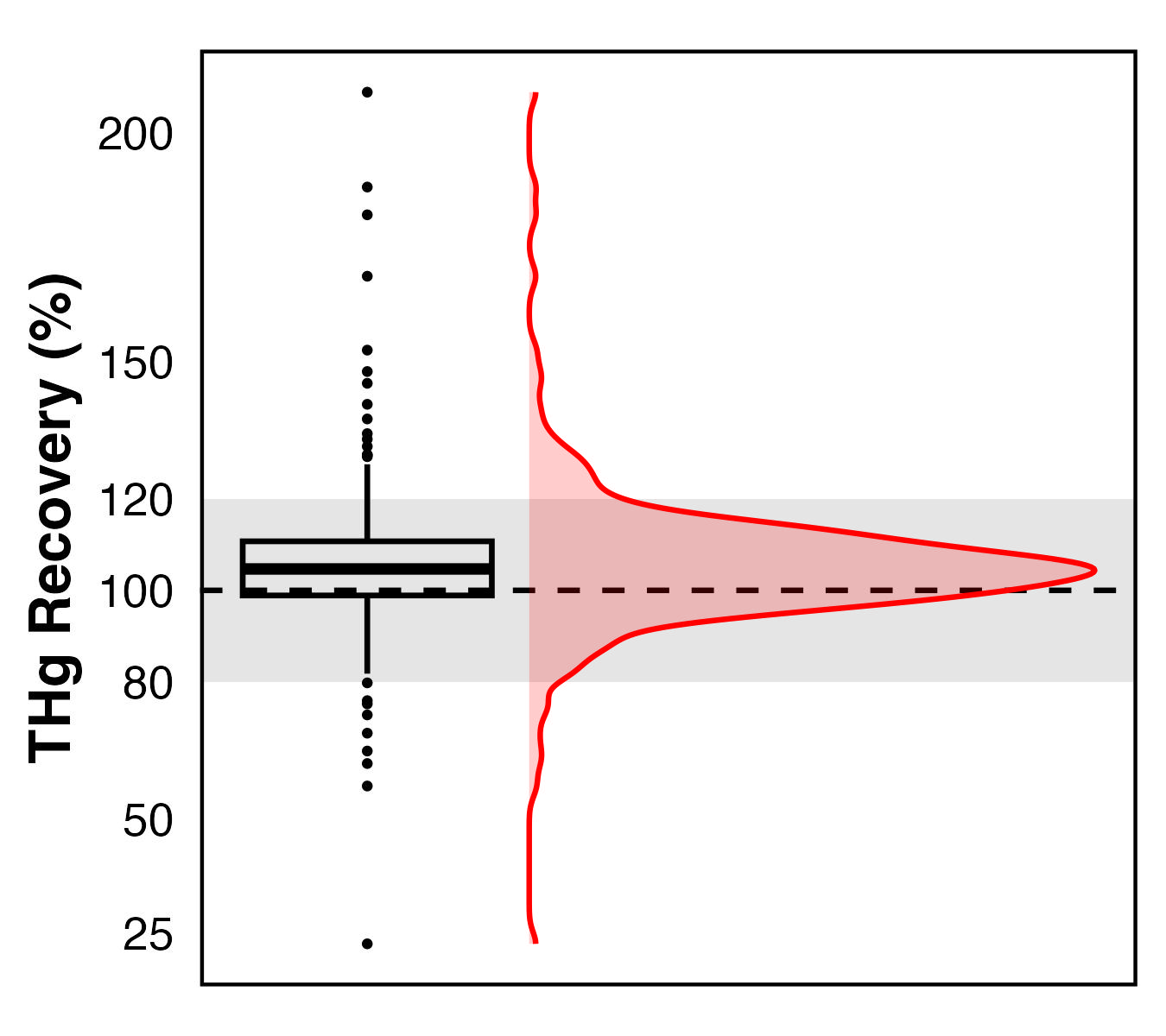

### FigS5_Recovery_Heteroscadasticity.jpg

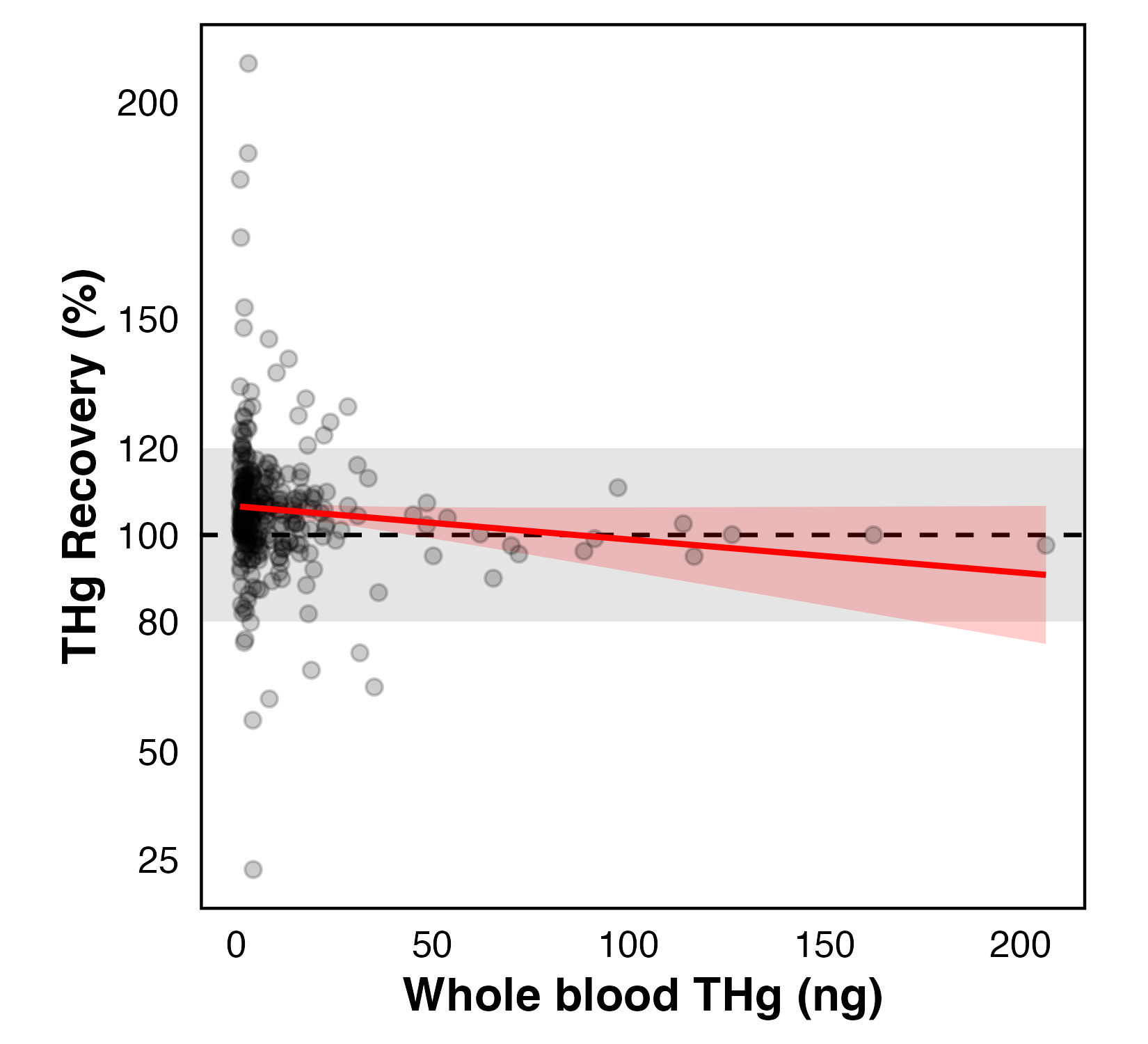
